# Genetic profiling via a novel PCR-RFLP method enabled identification of four genera of anaerobic gut fungi isolated from nyala, giraffe, and zebra hosts

**DOI:** 10.64898/2026.05.25.727589

**Authors:** Louisa Edge, Peiru Duan, Stephane Kerangart, Alice M. Buckner, Jolanda M. van Munster

**Affiliations:** School of Natural and Social Sciences, Scotland’s Rural College, Edinburgh, UK

**Keywords:** fungal diversity, herbivore digestive system, rRNA gene PCR-RFLP, *Feramyces*, *Neocallimastix*, *Khoyollomyces*, *Piromyces*, NY08, new species

## Abstract

Herbivore gut microbiomes may contain a diversity of anaerobic gut fungi (AGF, phylum *Neocallimastigomycota*), important for fibre degradation. To perform functional studies and elucidate niches of different AGF species, representative fungal isolates must be obtained into axenic culture, which is a resource-intensive process. Here we leverage the integration of morphological and functional assessments of AGF isolates with a newly developed PCR-RFLP strategy, to distinguish and identify isolates of interest from faecal samples from zoo-housed animals. In silico prediction of PCR-RFLP profiles of cultured genera, followed by experimental validation, confirmed that LSU-based PCR-RFLP with AluI and Hyp188I digestion was effective in identification of fungi of distinct genera. Together our workflow resulted in isolation of a so far uncultured *Piromyces* (NY08) species and *Neocallimastix cameroonii* from nyala samples, as well as *Feramyces austinii* from giraffe and *Khoyollomyces ramosus* from zebra. Amplicon sequencing confirmed that these species dominated AGF communities in their hosts, likely benefiting isolation success, and we identified enrichment conditions which also affected cultivability. The workflow developed here aids efficient AGF isolations, which are instrumental in expanding opportunities for functional studies that provide insight into the physiology and ecology of these fungi and help realise applications in white and green biotechnology.

**One sentence summary:** A validated PCR-RFLP strategy, developed based on genetic diversity data from *Neocallimastigomycota*, enables efficient identification of isolates of these anaerobic gut fungi from environmental samples, as demonstrated via targeted enrichment of anaerobic gut fungi common in faeces of giraffe, zebra and nyala, resulting in isolation of species of genera *Feramyces*, *Neocallimastix*, *Khoyollomyces,* and a novel *Piromyces/*NY08 species.

## Introduction

The digestive tract of ruminants contains a microbiome that enables conversion of plant material to nutrients for its host. This digestion of feed is critical to ruminant health, and to their performance in farming. Driven by this agricultural importance, major advances have been made in understanding the diversity and function of bacterial and archaeal microbiome members (Mizrahi et al., 2021; Seshadri et al., 2018; Solden et al., 2018), but eukaryote members, anaerobic gut fungi (AGF, phylum *Neocallimastigomycota*) and protozoa, are less well studied, despite their importance in digestion. AGF activity increases feed digestion, weight gain and milk production *in vivo* in ruminants, as reviewed (Gruninger et al., 2014), and increases digestibility of feed by 13-53% in rumen fluid *in vitro* cultures (Lee et al., 2000; Ma et al., 2020). The effective AGF digestive activity is thought to be enabled by a broad arsenal of digestive carbohydrate active enzymes as well as penetrative fungal structures that open up plant material for further digestion (Akin & Borneman, 1990; Haitjema et al., 2017; Solomon et al., 2016).

Genomics-based approaches have revealed a wealth of AGF diversity, but isolation, cultivation, and functional characterisation of AGF is falling behind. Amplicon sequencing using the ITS region or 28S rRNA (LSU) gene markers uncovered the existence of a broad variety of AGF genera in ruminants and hindgut fermenters (Edwards et al., 2017; Kittelmann et al., 2012; Liggenstoffer et al., 2010; Meili et al., 2023; Nicholson et al., 2010). More recently, AGF were also found to be present more broadly in herbivores, including ostrich (Vinzelj et al., 2025) and tortoises (Pratt et al., 2024). Obtaining cultured isolates of these fungi allows for direct functional studies of properties such as metabolism or lignocellulose degradation capacity, and exploration of their capabilities for biotechnology applications. Isolation efforts over the last decade have resulted in the cultivation of more than a dozen new fungal genera from the digestive tracts of domestic and wild herbivores (Dagar et al., 2015; Hanafy, Johnson, et al., 2020; Hanafy, Lanjekar, et al., 2020; Pratt et al., 2024; Vinzelj et al., 2025), as reviewed by (Hanafy et al., 2022). However, there are numerous uncultivated anaerobic fungal lineages in both wild and domesticated herbivores (Meili et al., 2024), and for many AGF genera currently only one species has so far been brought into culture (Hanafy et al., 2022). Evidently, a substantial diversity of AGF exists in nature that has so far not been cultured. Furthermore, considerable challenges exist in long-term preservation and transport of AGF isolates (Elshahed et al., 2022; Nagpal et al., 2012; Vinzelj et al., 2022). Together, this results in a limited availability of a diversity of AGF strains, hampering functional studies of AGF.

With increasing interest in the AGF for their biotechnological and agricultural applications, and engagement of a growing global community of researchers with this field, it is of increasing importance to report on, and ensure the availability, of broadly accessible strategies to obtain isolates of AGF into culture and assess their identity. In bringing AGF into culture, the diversity of fungi isolated from a sample is thought to be affected by interplay between the diversity and evenness in the sample’s microbiome as well as the employed enrichment and isolation conditions. AGF are found along the ruminant digestive tract (Davies et al., 1993). While AGF communities in faecal samples are less diverse and distinct in composition than in rumen-derived samples, as summarised in (Meili et al., 2024), faecal samples are far more easily obtained for a wide range of animals. AGF enrichment from stool samples, followed by isolation of fungal strains, is therefore commonly practised in the field and has resulted in recovery of a range of novel genera. While some herbivores harbour a very diverse AGF community in such samples, other host AGF communities seem to be dominated by one or few species (Meili et al., 2023). A high relative abundance of an AGF species in such samples, and a low evenness of the AGF community, has been demonstrated to strongly correlate to successful isolation of the AGF (Hanafy, Johnson, et al., 2020), as also suggested by Stabel et al (2020). Also, the composition of culture media has been reported to affect which AGF are enriched, for example the carbon source used affects diversity (Griffith et al., 2009) and the inclusion of rumen fluid selectively enriches for *Neocallimastix* and *Piromyces* genera (Joshi et al., 2022).

Selection or characterisation of any obtained AGF isolates may include microscopy or functional testing, but unambiguous identification requires DNA-based analysis as key morphological properties such as number of zoospore flagella (mono or poly-flagellate), thallus development (mono or polycentric), or rhizoid morphology (bulbous or filamentous) are commonly shared by taxa, and other distinctive morphological characteristics are subtle and be easy to miss in initial non-axenic isolate cultures (Hanafy et al., 2022). DNA-based methods such as Sanger sequencing of ribosomal ITS and 28S LSU regions or amplicon sequencing are commonly employed (e.g. (Hanafy, Lanjekar, et al., 2020; Joshi et al., 2022; Wang et al., 2017; Young et al., 2022), but these may be costly in terms of time or reagents. Historically, DNA-based assessments of the ITS or 28S LSU region via PCR-DGGE or -RFLP has been used to delineate similar groups of isolates, as reviewed by (Edwards et al., 2017), methods that may be fast and economical but were not able to confirm isolate identity. However, their use for discrimination between *Orpinomyces* species (Dagar et al., 2011) and between up to six distinct genera has been demonstrated (Dagar et al., 2014; Fliegerová et al., 2006; Hausner et al., 2011), suggesting potential for developing this technique via integration with genetic information of the broad range of AGF genera that has become available over the last decade.

In this study we aimed to isolate a set of AGF species representing different genera and test a combination of identification methods to facilitate fast and economical isolate identification. We hypothesized that integrating different functional and molecular characterisation methods (Fig.1) would benefit assessment of the diversity of culturable AGF isolates obtained from faecal samples. Zoo-housed herbivores giraffe, nyala and zebra were selected as hosts as these animals represent different gut systems, with nyala and giraffe being ruminants and zebra a hindgut fermenter. The zebra AGF community has been reported to be rich in *Khoyollomyces* and the giraffe’s in *Feramyces*, with faecal samples of these animals therefore likely to harbour distinct AGF (Hooker et al., 2023; Meili et al., 2023). The nyala mycobiome has to our knowledge not been investigated so far. From these samples we successfully obtained AGF isolates representing *Neocallimastix*, *Feramyces*, *Khoyollomyces* and *Piromyces*/NY08 genera. Isolation success was critically affected by vitamin and antibiotic supplementation of the cultivation media. Based on genetic information of the D1/D2 LSU region of 21 established AGF genera, we developed and validated a PCR-RFLP method that was instrumental in time and cost-effective identification of fungal isolates.

**Figure 1.**
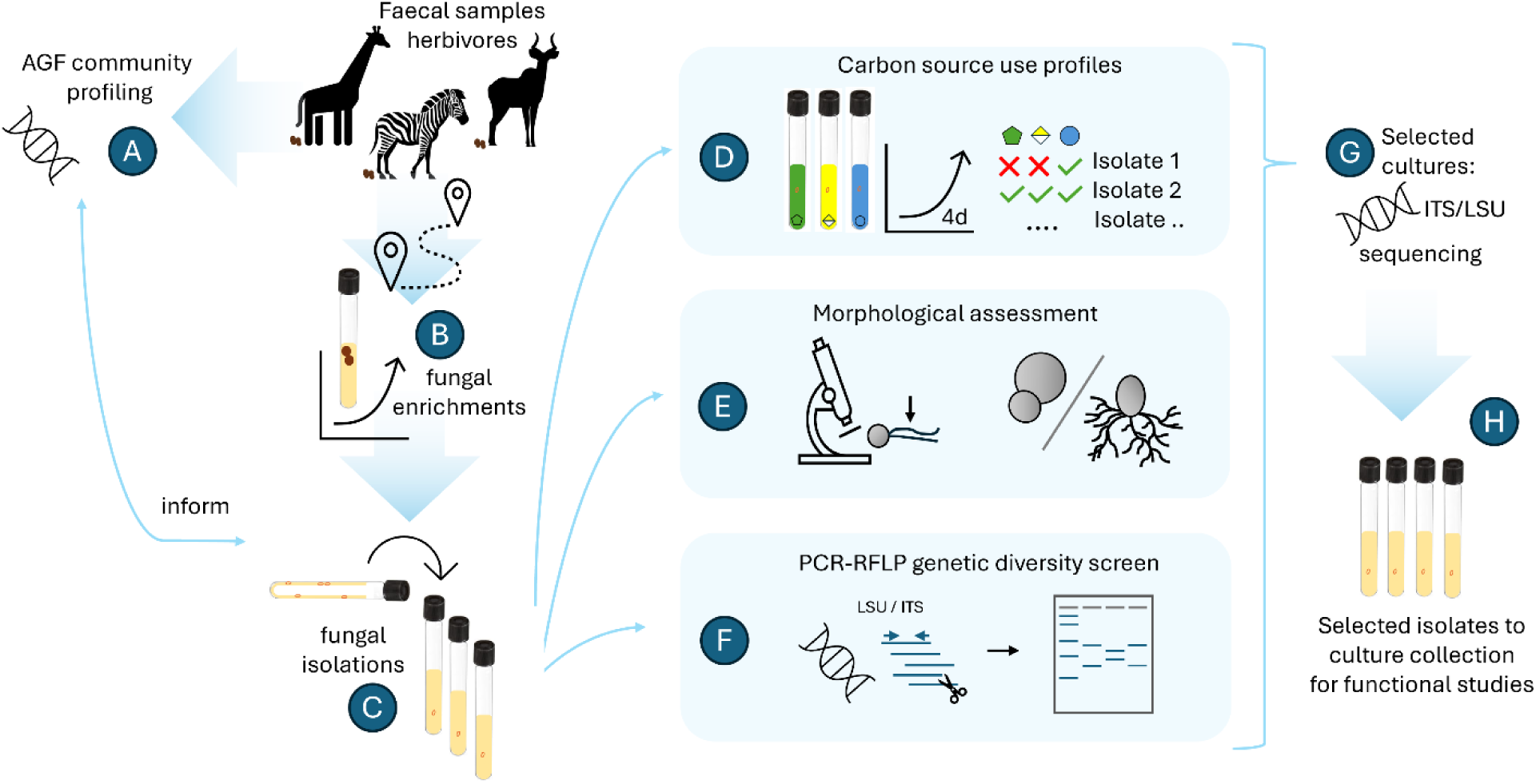
Approach for isolation and characterization of a diversity of AGF. **(A)** AGF community profiling via amplicon sequencing informs isolations, **(B)** faecal samples from wild ruminants kept in captivity, giraffe and nyala, used as inoculum for enrichment cultures, from which **(C)** isolations took place via roll tube cultivation. Isolates were characterized via **(D)** measurements of growth on fructose, galacturonic acid and glucose as carbon source, **(E)** assessment of morphology and zoospore number, **(F)** PCR-RFLP of ITS or LSU region to profile genetic diversity, followed by **(G)** confirmation of identities for selected isolates via ITS and/or LSU rRNA gene sequencing and **(H)** further selection for functional studies.

## Materials and Methods

### Faecal samples

Faecal samples were obtained from nyala antelope (*Tragelaphus angasii*), giraffe (*Giraffa camelopardalis*), and Grevy’s zebra (*Equus grevyi*) housed in Edinburgh Zoo, Scotland, United Kingdom. Samples were collected freshly after deposition, by trained zoo personnel following protocols for entry to animal enclosures approved by Edinburgh Zoo. Faecal samples were transported to the laboratory and stored at -20°C until use.

### Fungal enrichment, isolation and growth

Fungal cultivations and enrichments, unless specified otherwise, were performed in 6 ml medium C (as described by (Theodorou et al., 2005) but replacing bactocasitone with tryptone) in 15 ml Hungate tubes containing a CO_2_ atmosphere and incubated at 39 °C for 3-4 days.

For initial AGF enrichments, faecal material was dispersed in 6 ml medium C and subsequently diluted 20-fold into 6 ml medium C, with 1 % w/v 0.5 mm wheat straw particles. Cultures were supplemented with antibiotics to repress bacterial growth as follows: 50 µg ml^-1^ chloramphenicol (for giraffe samples) or 100 µg ml^-1^ penicillin and 100 µg ml^-1^ streptomycin (for nyala samples). After 4 days, when floating plant material and gas pressure accumulation indicated possible fungal activity, aliquots from the enrichment cultures were used to inoculate roll tubes consisting of 2 ml medium C with 5 g l^-1^ glucose and 15 g l^-1^ agar, supplemented with the corresponding antibiotics, in 15 ml Hungate tubes. After incubation for 3 days at 39 °C, formation of fungal colonies was confirmed via microscopy. Then, 20 isolated fungal colonies were picked under anaerobic atmosphere (80% N, 10% H_2_, 10% CO_2_) in a Whitley A25 workstation (Don Whitley Scientific) and transferred to tubes with 3 ml medium C with 1 % w/v wheat straw and 50 µg ml^-1^ chloramphenicol. Obtained cultures were maintained by transfer of 0.3 ml to a new tube of culture medium every 3-4 days.

Additional enrichments for nyala samples were performed after 8 months of sample storage at -20°C, as described above, except replacing chloramphenicol with 100 µg ml^-1^ penicillin and 100 µg ml^-1^ streptomycin during culture maintenance. For additional enrichments from zebra samples, a panel of antibiotics and vitamins was trialled, including combinations of 50 µg ml^-1^ chloramphenicol, or 100 µg ml^-1^ penicillin and 100 µg ml^-1^ streptomycin, 50 µg ml^-1^ kanamycin, as well as the chloramphenicol or ampicillin/streptomycin together with 100 µl vitamin solution 2. Vitamin solution 2, described in (Teunissen et al., 1991), consisted of 5 mg l^-1^ thiamin.HCl, 5 mg l^-1^ riboflavin, 5 mg l^-1^ calcium D-pantothenate, 5 mg l^-1^ nicotinic acid, 2 mg l^-1^ folic acid, 1 mg l^-1^ cyanocobalamin, 1 mg l^-1^ biotin, 10 mg l^-1^ pyridoxin.HCl and 5 mg l^-1^ p-aminobenzoic acid.

### Growth profiles for carbon source use

The ability of fungal isolates to use different sugars as carbon source was screened in medium C containing either 5 g l^-1^ glucose, fructose, or galacturonic acid as the sole carbon source. Sugars were added to the autoclaved media as filter-sterilised stock solution and 50 µg ml^-1^ chloramphenicol was supplied to repress bacterial growth. Fungal isolates were initially cultured on medium C with wheat straw as carbon source for 3 days to achieve an exponentially growing culture, after which a 10% v/v inoculum was transferred to initiate cultures for profiling of carbon source use. Growth was assessed after 4 days of cultivation, via visual assessment of fungal biomass formation, and measurements of fermentation gas pressure. For selected isolates, additional rounds of purification were performed to achieve axenic cultures, after which C-source use profiles were confirmed via cultivation in the media described above.

### DNA extraction

Fungal isolates were grown in 6 ml or 50 ml cultures with glucose as carbon source for 3 – 4 days, after which biomass was harvested via centrifugation (7 min at 3893 *g*), frozen and lyophilized. DNA was extracted using a CTAB extraction of pulverised biomass followed by chloroform extraction and isopropanol precipitation, as described (Griffith et al., 2009) but replacing the hot ethanol wash step with ambient ethanol. Obtained DNA was used to perform PCR-RFLP or ITS2-based amplicon sequencing.

For DNA from stool samples sent for AGF-specific amplicon sequencing with LSU-EnvS primers, DNA was extracted with the Qiagen DNeasy Plant Pro Kit with the following modification: cells in stool samples (∼ 100 mg wet weight) in CD1 lysis buffer were disrupted for 3 x 1 min in a Qiagen Tissuelyser LT at 50 Hz, followed by a 10 min incubation at 65°C, before continuing purification as described by the extraction kit manufacturer.

### PCR-RFLP

PCR-RFLP using the ITS region was performed generally as described by (Fliegerová et al., 2006). In short, PCR was performed with primers ITS1 and ITS4 using Promega GoTaq Mastermix, whereafter product formation was confirmed via agarose gel electrophoresis. PCR products were purified from 20 µl PCR mixture using the Wizard® SV Gel and PCR Clean-Up System (Promega) as directed by the manufacturers. PCR products were eluted in water and ∼ 0.5 µg DNA was digested with 5U of DraI or HinfI at 37°C for 1 h, after which the resulting DNA fragments were visualised on a 2.5% agarose gel using Gelred nucleic acid stain and ThermoScientific GeneRuler 100 bp ladder. PCR-RFLP using the LSU region was performed similarly, but using the primers NL1 and NL4, and digestion with Hpy188I or AluI, followed by visualisation on a 3% agarose gel.

To generate an ITS PCR-RFLP database of predicted PCR-RFLP fragments for each of the so far identified species of AGF, the AF_Full_Regiondatabase version 2.0 was obtained from the Anaerobic Fungi Network (www.anaerobicfungi.org). Initial sequence alignments were performed in Clustal Omega (Sievers et al., 2011). Using R 4.4.2 with the R-packages DECIPHER version 3.2.0 (Wright, 2024) and Biostrings version 2.74.1 (Pagès et al., 2025), the sequences were trimmed to the ITS1 and ITS4 primer borders and complete sequences were separated from incomplete sequences (lacking the ∼ 50 bp at the ITS1 border). Following in silico digestion with DraI and HinfI restriction enzymes, fragment sizes were tabulated. For selected sequences, results were visualised on an in silico 2.5% agarose gel using SnapGene 8.2.1. (Fig 3). For Fig. 3A, DraI digestion, the following sequences were used: JQ782543, JQ782547, MT085722, Aoud18_14398, MT085694, MW907587, JQ782554, MZ044643, MT085707, MW899532, MT085733, JQ782551, JQ782553, MT085711, MT085666, MT0856676, MT085680, MT085712, MT085689, Horse_10178, AxDoe_98435, ZebrA149764, MT085698, MT085728, MT085727, MT085708, MT085709, MT085738, OQ382931 and OQ382915. For Fig. 3B sequences were as for DraI digestion except, to display the most representative profiles, JQ782547, MZ044643 and MT085709 were omitted, MT085707 was replaced by MT085705, MT085733 by MG605690, JQ782551 and JQ782553 by MT085734, and ZebrA146613, MG605690 were added.

For development of an LSU-based PCR-RFLP, the AF_LSU_database version 2.0 was obtained from the Anaerobic Fungi Network (www.anaerobicfungi.org) and sequences were trimmed to NL1 and NL4 primer sequence borders. Sequences lacking the NL4 primer sequence were separated from complete sequences and in silico digestion with a range of restriction enzymes was tested for ability to generate fragments that allowed discrimination between AGF genera and/or species. For selected sequences, results were tabulated and visualised on an in silico 3% agarose gel using SnapGene 8.2.1. The following sequences are used for visualisation in Fig. 5: for AluI: JQ782543, MT085722, MW019480, MT085694, MW907586, JQ782554, MT085707, MW899532, MG605686, MT085733, MT085711, MT085668, MT085675, MT085679, Piromyces_spUH3_1_LSU_scaffold317_9682_10448, KP205570, Zebra_149764, MT085698, MT085701, MT085728, MT085709, MT085738, OP253962, OP253957. For Hpy188I, to display the most representative profiles, MW907587 replaced MW907586, MT085666 replaced MT085668 and Horse_101214_Piromyces was added. Predicted LSU PCR-RFLP profiles were subsequently experimentally validated as described above using DNA obtained from a panel of AGF strains available in the laboratory: *N. frontalis* CoB3 (Shen et al., 2026), *N. cameroonii* N9 and N13, *Caecomyces communis* SHB (Shen et al., 2026), *Capellomyces forminis* Cap2A (Hanafy et al., 2023), *Piromyces edwardsiae* isolate SHC (Shen et al., 2026) and *Feramyces* sp. isolates G3 and G5.

### Amplicon sequencing

DNA of isolates and faecal samples was sent to Novogene for amplicon sequencing of the ITS2 rRNA gene with primers ITS3-2024F (5’-GCATCGATGAAGAACGCAGC-‘3) and ITS4-2409R (5’-TCCTCCGCTTATTGATATGC-‘3), or for 28S LSU rRNA gene region with primers AGF-LSU-EnvS-F (5’-GCGTTTRRCACCASTGTTGTT-‘3) and AGF-LSU-EnvS-R (5’-GTCAACATCCTAAGYGTAGGTA -‘3) (Young et al., 2022) that produce a ∼ 370 bp fragment, using amplification conditions as described in (Meili et al., 2023). The negative controls did not yield any amplification products and were omitted from further analysis. Illumina 250 bp paired end sequencing was used via the NovaSeq 6000 to obtain a minimum of 30k raw reads per sample. Sequences were analysed using Mothur v.1.48.5 (Schloss et al., 2009) following a protocol based on (Meili et al., 2023) and (Kozich et al., 2013). Demultiplexed reads were assembled to contigs while removing primer and barcode sequences. Quality trimming was performed to include only sequences between 300 and 320 bp in length, with homopolymer stretches of ≤ 8 bases, that did not contain ambiguous bases and aligned to *Neocallimastigomycota*. Preclustering of the 19.8-64k reads per sample was performed allowing 3 nucleotide differences and chimeras were removed via the chinera.vsearch command using dereplication. Sequences were classified using the aforementioned AF_LSU_database (v2), and OTUs were generated using a cutoff of 0.03 and classified using the same database. Metadata, taxonomy and OTU tables were imported into the R-package phyloseq (McMurdie & Holmes, 2013) for analysis and data visualisation. Alpha-diversity (Observed, Shannon, and Simpson Indices) was calculated from raw counts and tested for significance using a Kruskal-Wallis chi-squared test followed by Dunn’s Pairwise Comparison with Benjamin-Hochberg correction for multiple testing. Taxa were pruned to include only those with mean > 0.5 reads, resulting in 169 OTUs across 9 samples. Data were converted to relative abundances, beta-diversity calculated via weighted UniFrac ordination and PCoA analysis, and tested for significance using PERMANOVA via adonis2 in the vegan package (Oksanen et al., 2026).

## Results

Faecal samples of zoo-housed zebra, nyala and giraffe were used for amplicon-based profiling of resident AGF communities (Fig. 1A), and also as source for isolation of AGF strains (Fig 1B, C), which we characterised using a combination of three methods (Fig. 1D-F).

### Zebra and nyala, but not giraffe harbour simple AGF communities

To assess AGF diversity in the host mycobiome, we performed amplicon sequencing using AGF specific primers AGF-LSU-EnvS (Young et al., 2022). After filtering 169 OTUs were obtained over the different samples. Taxonomy (Fig. 2A,B) indicated that the AGF community from zebra consisted almost exclusively of (99.8-99.9% of reads) *Khoyollomyces*. The AGF community from nyala also had limited α-diversity and mainly consisted of *Neocallimastix cameroonii* and an uncultured proposed genus NY08 that is closely associated (Meili et al., 2023) with *Piromyces*. The giraffe faeces harboured an AGF community with higher α-diversity, in terms of Observed, Shannon as well as Simpson indexes (Fig. 2C), with 3 distinct dominant genera (Fig. 2A). Besides *Neocallimastix frontalis*, and *Feramyces austinii*, the giraffe samples were also abundant in *Piromyces*-associated NY08 (100% identity of respective OTU sequences). In addition, up to 4.8% of *Neocallimastigomycota* reads in giraffe sequences could not be classified to genus level. Notably, while the proposed genus NY08 previously was only identified to have <5% abundance over a range of animal samples (Meili et al., 2023), it represents 6-44% of reads in the giraffe samples and 46-94% of reads in the nyala samples, suggesting that these samples may be a good source to isolate this so far uncultured fungus from. Based on the PCoA analysis the three animals harboured significantly different communities (*p*=0.007), with PC1 and PC2 explaining 32.9% and 24.7% of the variation observed, respectively. Distinct clustering was observed by host animal (Fig. 2D), however the samples did not seem to cluster by gut type.

**Figure 2.**
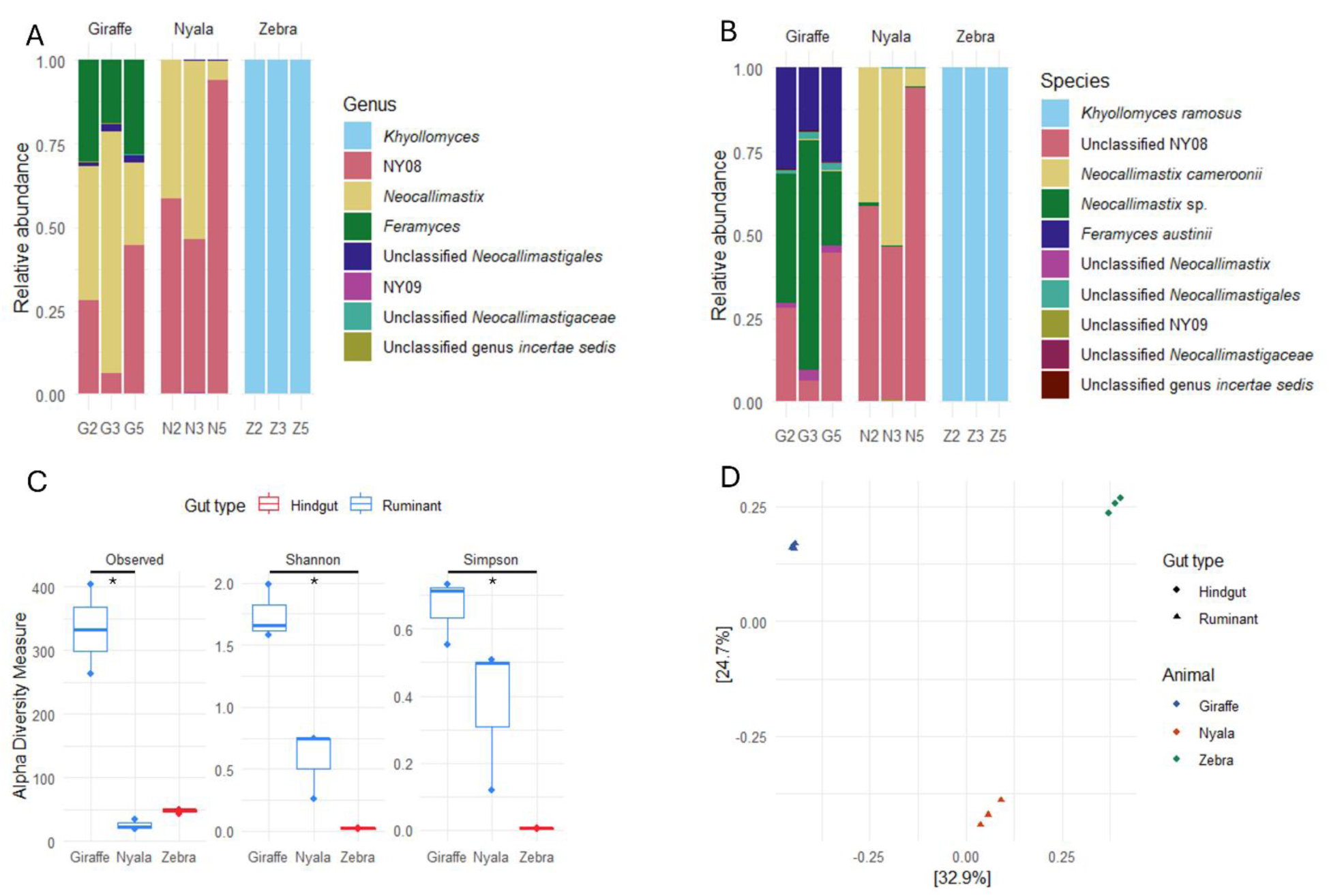
AGF community composition in original stool samples. Amplicon sequencing, with AGF-specific primer pair AGF-envS F/R, of stool samples with **(A)** genus assignment, **(B)** species assignment **(C)** α-diversity (Observed, Shannon, and Simpson) with significance tested via a Kruskal-Wallis test followed by Dunn’s Pairwise Comparison with Benjamin-Hochberg correction. *ns* indicates *p-*value > 0.05, * a *p*-value of [0.05 – 0.01).

### Obtaining fungal isolates from faecal samples

Stool samples were used to inoculate AGF enrichment cultures in complex medium C with wheat straw as carbon source, a complex fibrous plant material that can be degraded by broad range of AGF (Fig. 1B). The obtained enrichment cultures were used to inoculate solid medium in roll tubes (Fig. 1C). Roll tubes inoculated from enrichments originating from zebra samples did not initially yield any colonies, despite multiple attempts to repeat enrichments and roll tube cultivation, employing different combinations of antibiotics.

For roll tubes inoculated from enrichments from nyala and giraffe samples, twenty colonies were subsequently picked per enrichment, of which in total 80% were successfully taken through all (27 isolates) or a subset (5 isolates) of the characterisation steps, with the remainder lost from cultivation. Isolates were characterised, via a combination of profiling of carbon source use, morphological assessment and molecular biology (PCR-RFLP and sequencing) (Fig. 1D-F), to assess diversity of cultured isolates and to prioritise isolates for identification and further purification to the axenic cultures required for full characterisation and functional studies. Efficacy of the three characterisation techniques, in terms of reliability and resource use, were also assessed.

### Profiling of carbon source use of isolates from nyala

Growth on plant-derived simple sugars fructose, glucose and galacturonic acid was assessed, whereby glucose and fructose are expected to be metabolised by all or most AGF genera and species (Trinci et al., 1994). The plant cell wall polysaccharide pectin, or its monomer galacturonic acid, is not commonly metabolised by AGF, with exception of *Feramyces austinii* (Hanafy et al., 2018), and some *Neocallimastix* isolates (Kopečný & Hodrová, 1995). We recorded fungal growth via fermentation gas levels, a standard in the field used as a quantitative proxy for growth of AGF (Theodorou et al., 1995; Wilken et al., 2020) as well as visual observations of fungal biomass. However, we found considerable discrepancies between both (Fig. S1) with false positives, possibly due to bacterial contamination of the initial fungal isolates, as well as some false negatives, which indicated that gas pressure measurements alone are ineffective when assessing growth of non-axenic fungal isolates in our workflow.

Results of visual observation of generated fungal biomass (Fig. 3A) demonstrate that after 4 days all isolates used glucose and fructose to support their growth, and most used galacturonic acid, except isolates N4, N5, N6 and N13. This indicated that there seem to be two distinct carbon source use profiles between AGF isolates from nyala, suggesting metabolically diverse strains have been isolated. To confirm, representative isolates, N2, N6, N9, and N13 were purified to axenic cultures, and used to generate growth profiles (n=3) over 7 days. However, while all strains grew well on glucose and fructose (Fig. S2A) generating 10.7-18.1 mg dry weight of biomass per culture, on galacturonic acid the accumulated gas pressure accumulation was negligible and biomass only 2.0-5.4 mg per culture. Overall, this suggests that growth of all these isolates on galacturonic acid was slow or limited, and initial observed differences during growth on galacturonic acid were likely reflections of variations of growth rather than differences in innate capability.

**Figure 3.**
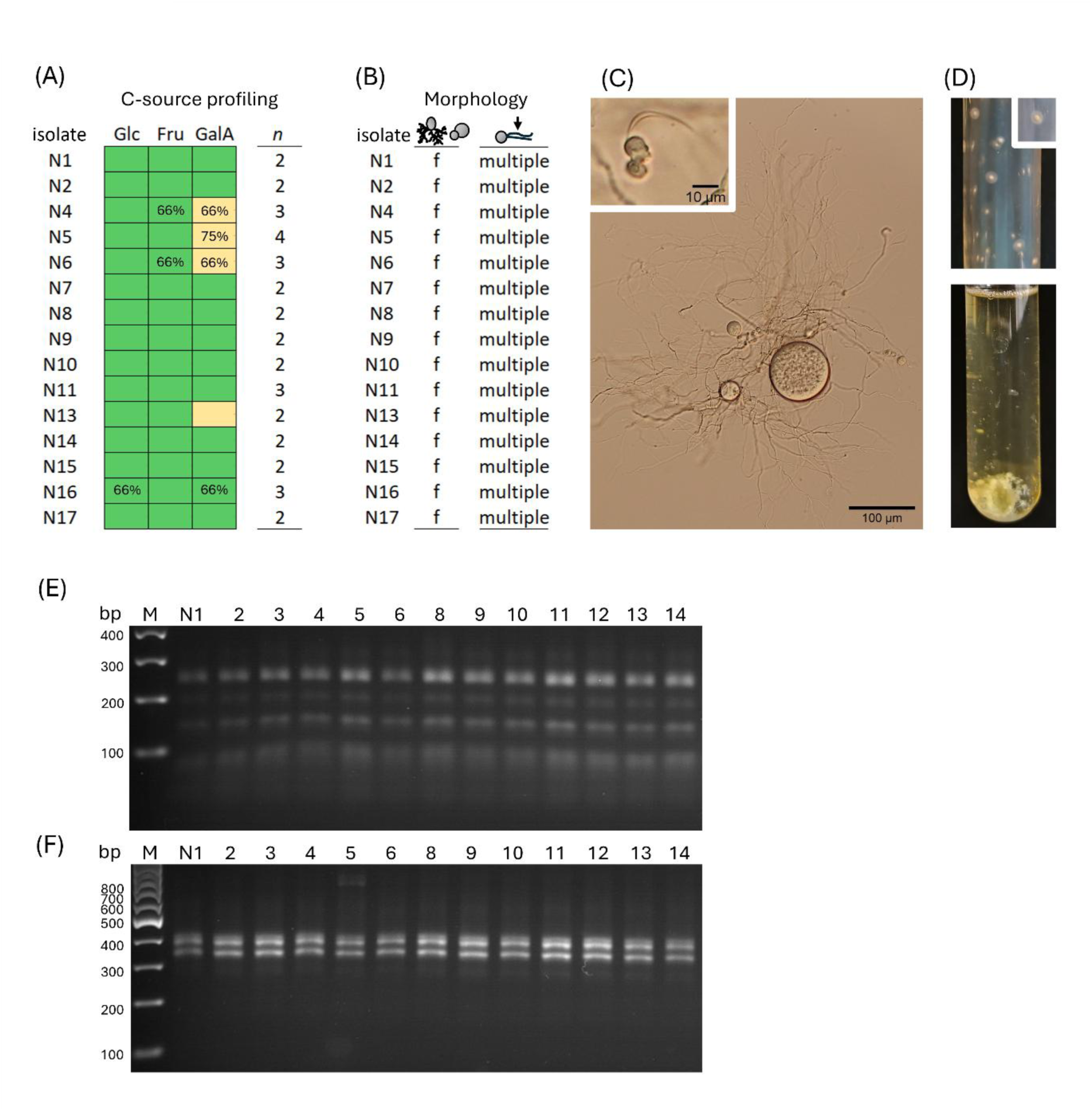
Characterisation of AGF isolates obtained from nyala. **(A)** Observations of fungal biomass formed in cultures with glucose (Glc), fructose (Fru) or galacturonic acid (GalA) as carbon source, green indicating biomass formation, yellow no biomass. Data given as most representative of replicate cultures, with *n* indicating the number of replicates, conditions with percentages indicating the number of replicates matching the results shown, conditions without percentages gave identical results (100%). **(B)** morphology classification for each isolate, morphology as filamentous (f) or bulbous (b), number of flagella on zoospores indicated as single or multiple, **(C)** representative morphology of isolates (N9), zoospore with flagella visible (inset), **(D)** representative macromorphology, of isolate N9 grown in liquid media C with glucose as carbon source for 3d, and on solid media as roll tube for 2d and (inset) 10 d. **(E)** PCR-RFLP of selected isolates with ITS rRNA gene region digested with DraI or **(F)** with HinfI.

### Morphological assessment of isolates from nyala

Microscopy identified isolates in liquid culture all had filamentous rhizoids and polyflagellated zoospores (Fig. 3B, C). The macromorphology of mycelium was similar in all cultures, whereby in initial growth stages biomass was visible in liquid media as dense mycelial balls that later merged in a sheer network of mycelium (Fig. 3D). On solid medium, small (2-3 mm) beige colonies formed that reached their maximum diameter in 2 days, with a dense centre that became dark brown over time. Following the key for AGF genera identification (Hanafy et al., 2022), morphology indicates *Ghazallomyces, Aestipascuomyces, Neocallimastix* may have been obtained.

### DNA-based profiling and identification of isolates from nyala as N. cameroonii

As morphological assessment was as expected insufficient for unambiguous identification, we combined this with a molecular approach. We initially aimed to use ITS-based PCR-RFLP to assess genetic diversity between isolates, as PCR of the ITS region followed by digestion with DraI and HinfI enzymes was successful in experimentally demonstrating discrimination between at least 2 AGF genera in RFLP (Fliegerová et al., 2006; Griffith et al., 2009). To assess if this approach was able to discriminate between the larger diversity of currently established genera and to enhance information derived from PCR-RFLP profiles, we integrated the PCR-RFLP with sequence information for ITS rRNA gene regions. Using the sequence database for the AGF genera described so far (AF_Full_Region database version 2.0 available from the anaerobic Fungi Network (www.anaerobicfungi.org)), sequence information corresponding to the regions amplified by ITS1 and ITS4 primers was extracted. While the ITS4 primer motif was identified in all database sequences, only 63 out of 197 sequences contained the ITS1 primer motif with remaining sequences having a 65-73 bp truncation at the 5’- end that created an uncertainty in the length of the first predicted RFLP fragment. In silico digestion with DraI or HinfI generated RFLP profile predictions for each cultured genus (Table S1), of which representative profiles were visualised (Fig. S3) to aid correlation with experimentally obtained RFLP profiles. Results of the in silico analysis indicate that most genera with cultured representatives can be distinguished using the predicted RFLP patterns. However, for some genera multiple RLFP patterns are predicted that likely reflect the previously reported diversity of ITS sequences within genera (Edwards et al., 2017).

Experimentally, amplification of the ITS region of our nyala-derived isolates with primers ITS1 and ITS4 resulted in a single PCR product. For all isolates DraI digestion generated 3 main bands (259 bp, 148 bp and 103 bp) and a faint additional band around 195 bp (Fig. 3E) that, based on intensity and cumulative size of all fragments, can only have arisen from partial digestion. HinfI digestion resulted in profile with 2 fragments, 392 and 348 bp in size (Fig. 3F). Repeat measurements of bands of known size indicate an error margin of 20-30 bp for absolute fragments sizes should be taken into consideration. When comparing reference patterns to the experimentally obtained profiles for nyala isolates (Fig. 2E-F), this indicated they correspond to *Neocallimastix cameroonii.* Specifically, experimental HinfI digestion patterns matched predictions for *N. cameroonii*, *Caecomyces, Capellomyces,* or *Liebetanzomyces*, while the DraI digestion results, taken uncertainty in the size of the first fragment into account, could match *N. cameroonii*, *Ghazellomyces,* and *Cyllamyces.* PCR-RFLP identification as *N. camerooni* is in line with the morphological characteristics identified for these isolates. Identification was further confirmed by ITS2 amplicon sequencing; sequences making up >99% of reads for isolates N2, N6, N9, and N13 had 96.1-99.7% identity to *N. cameroonii* clone 1 and 6 (Genbank MW175298, MW175302.1). Taken together, all fungal isolates obtained here from nyala consisted of *N. cameroonii*.

### Profiling of cultures from giraffe confirm isolation of Feramyces austinii

As our isolation efforts from the nyala stool sample yielded only one AGF species, despite initially picking 20 colonies, we repeated the analysis for a faecal sample from giraffe. Results of visual observation of fungal biomass in cultures on glucose, fructose and galacturonic acid (Fig. 4A), suggested that most isolates used all three carbon sources to support growth, whereas isolates G5 and G8 had not grown on galacturonic acid after 4 days. However, as observed for nyala isolates, verification with selected axenic isolates G2, G3, G5 over 7 days indicated that these strains eventually grew on all substrates (Fig. S2B), although with less biomass on galacturonic acid (3.4 - 5.9 mg dry weight per culture) compared to glucose and fructose cultures (12.7 - 17.9 mg per culture). Morphology of observed mycelium was similar for all isolates, with monocentric thalli, filamentous rhizoids and polyflagellated zoospores (Fig. 4B, C). In liquid culture, thick, dispersed growth on glucose and fructose was observed compared to more sheer growth on galacturonic acid (Fig. 4D). On solid medium light beige colonies with a dense, irregular shaped centre were formed that grew to over 5 mm diameter. These morphological characteristics correspond to those described (Hanafy et al., 2022) for genera *Ghazallomyces, Aestipascuomyces* and *Feramyces*.

**Figure 4.**
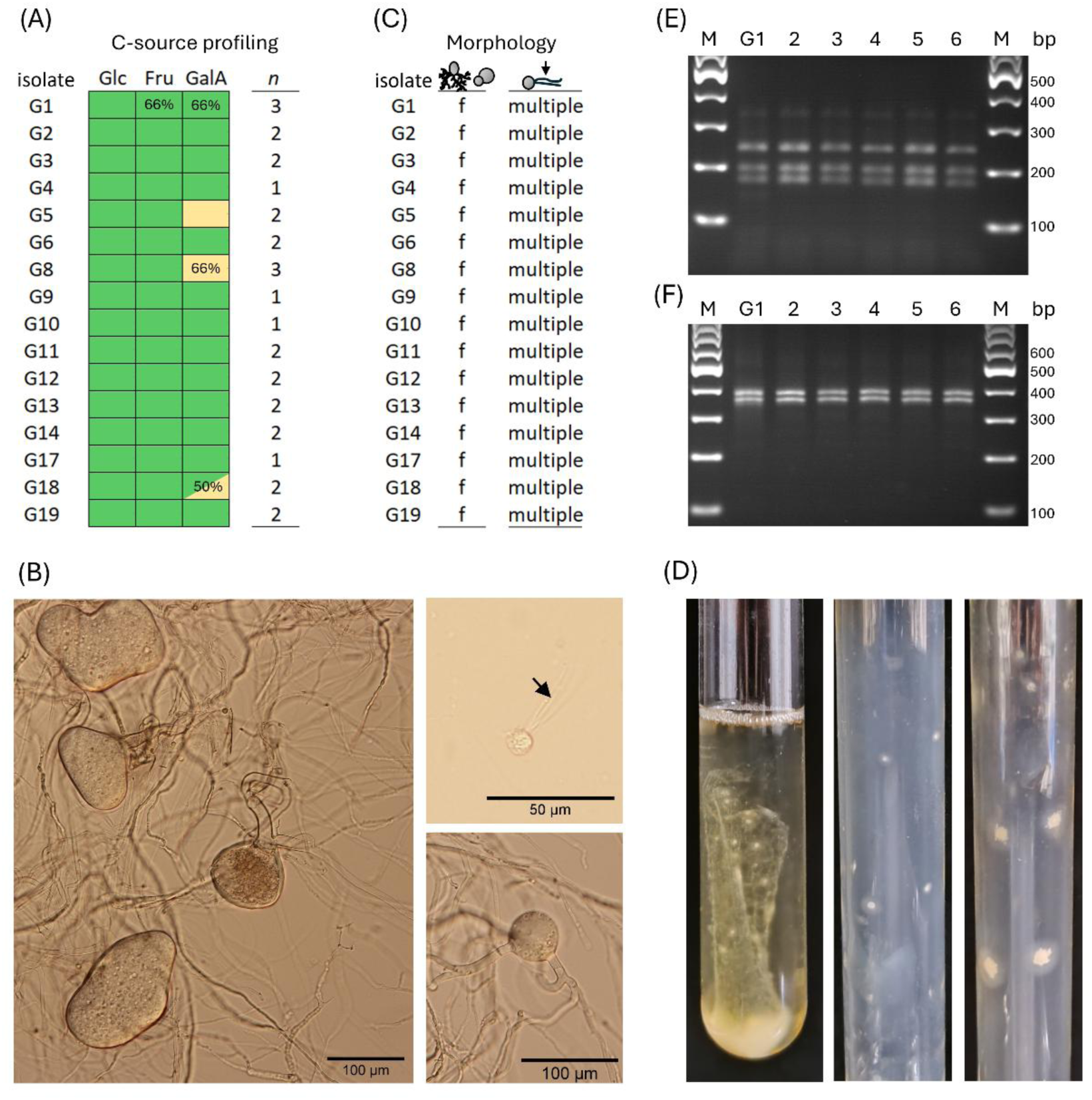
Characterisation of AGF isolates obtained from Giraffe. **(A)** Observations of fungal biomass formed in cultures with glucose (Glc), fructose (Fru) or galacturonic acid (GalA) as carbon source, green indicating biomass formation, yellow no biomass. Data given as most representative of replicate cultures, with *n* indicating the number of replicates, conditions with percentages indicating the number of replicates matching the results shown, conditions without percentages gave identical results (100%). **(B)** morphology of isolate G5 (left) and G2 (right), representative zoospore with flagella visible (arrow), **(C)** classification for each isolate, morphology as filamentous (f) or bulbous (b), number of flagella on zoospores indicated as single or multiple, **(D)** representative micromorphology, of 3 day old culture of isolate G3 grown in liquid media C with glucose as carbon source, and on solid media grown for 2 days (left) and 8 days (right). **(E)** PCR-RFLP of selected isolates with ITS rRNA gene region digested with DraI or **(F)** with HinfI.

PCR-RFLP resulted in a profile conserved among all isolates from giraffe but distinct from the nyala isolates. DraI digestion generated 3 distinguishable bands (Fig. 4E), determined to be 239, 191 and 166 bp, as consistent only with patterns predicted for *Feramyces*. HinfI digestion resulted in two visible fragments, measured as 390 and 363 bp (Fig. 4F), consistent with fragments predicted (Table S1) for *Feramyces, N. cameroonii, Ghazallomyces, Anaeromyces,* or *Capellomyces.* Together the PCR-RFLP indicated that all isolates likely belong to the genus *Feramyces*. This is consistent with our morphological observations and with the reported characteristics of *F. austinii* (Hanafy et al., 2018), the only known species in this genus. ITS2 amplicon sequencing resulted in >99% of reads from representative isolates G2, G3 and G5 having 98.4-100% identity to *F. austinii* strain DF1 and type strain F3a, thus confirming this identification.

### Improved PCR-RFLP strategy for identification for AGF using LSU sequences

ITS sequences have been reported to be variable and diverse in length between AGF genera and even demonstrate high diversity within species (Edwards et al., 2017). The ITS-based PCR-RFLP strategy described above has been useful for discrimination between fungal isolates but, as highlighted by Fig. S3 and Table S1, multiple common RFLP patterns were obtained for 6 out of 21 analysed genera and species for each of the restriction enzymes, with 4 patterns for *Piromyces.* This complicated tentative identification of isolates based on PCR-RFLP. As drawbacks of the ITS-based phylogenetic marker have resulted in a shift in the community towards using LSU-based identification (Elshahed et al., 2022; Young et al., 2022),and LSU-based PCR-RFLP enabled discrimination between *Orpinomyces joyonii* and *O. intercalaris* (Dagar et al., 2011), and between *Orpinomyces* and *Anaeromyces* isolates (Fliegerová et al., 2006), we explored if LSU-based PCR-RFLP could be used to aid fast identification of a wide range of AGF isolates.

We performed *in silico* digestion of LSU sequences with a range of restriction enzymes, assessing distinct fragment length polymorphisms between genus and species representatives. Digestion with the enzymes AluI (Fig. 5A) or Hpy188I (Fig. 5B) resulted in patterns of fragment lengths (Table S2) which, when combined, would enable differentiation between all genera. A single conserved or highly dominant pattern was found for all genera besides *Piromyces*. At the species level, *N. frontalis* and *N. cameroonii* can be easily distinguished on distinct RFLP, as can *O. joyonii* and *O. intercalaris*, while the RFLP for each of the two species represented in genera *Anaeromyces* and *Capellomyces* were identical or highly similar. The number of common patterns obtained for *Piromyces* was reduced compared to the ITS-based strategy but could not directly be linked to known species. This may reflect the phylogenetic complexity of the genus (Hanafy et al., 2023).

**Figure 5.**
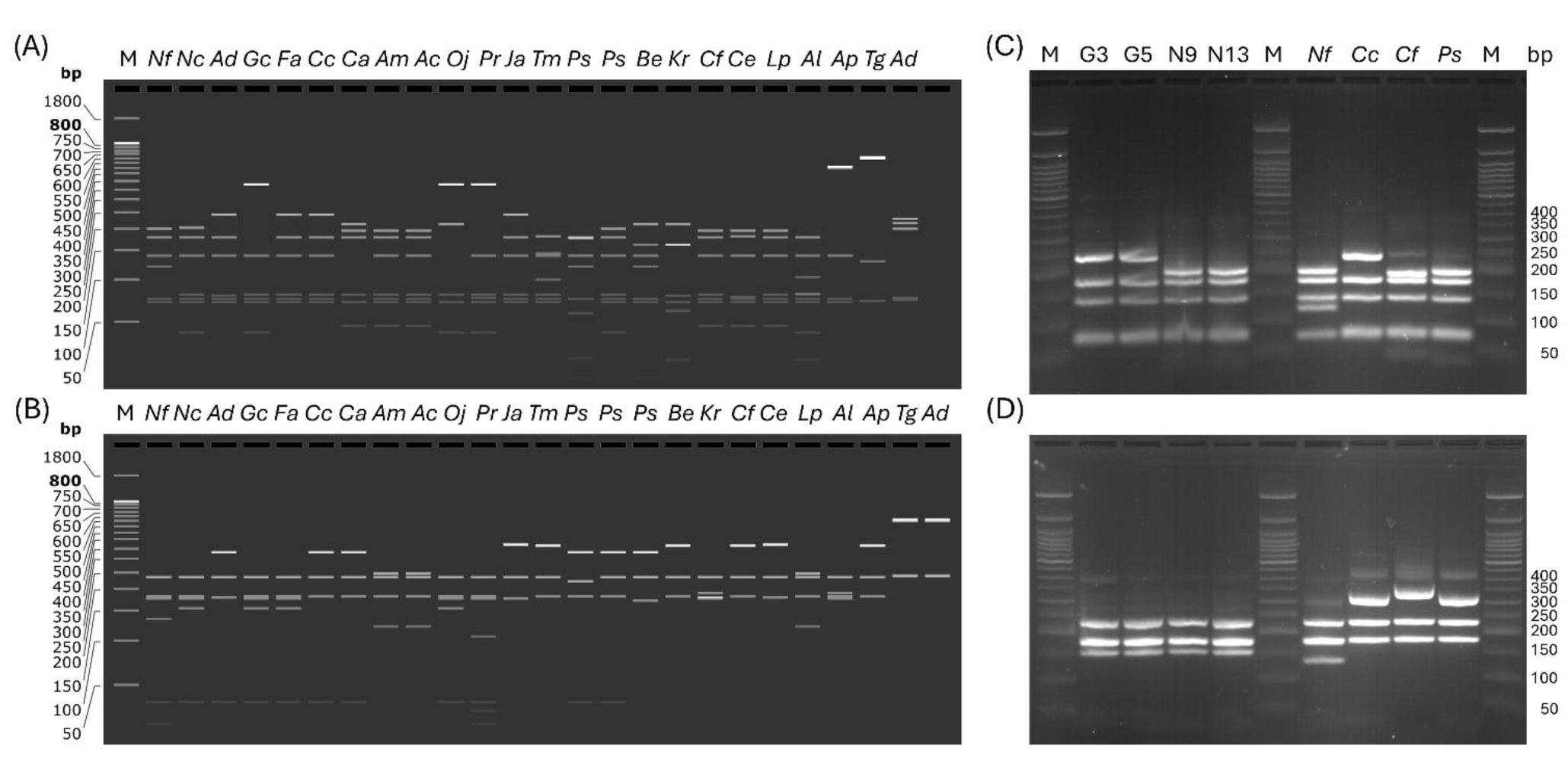
Theoretical and experimental validation of PCR-RFLP based on LSU sequence information. **(A)** In silico visualisation of RFLP profiles for selection of AGF genera and species predicted for digest of NL1-NL4 PCR product with AluI, and **(B)** with Hpy188I. *Nf Neocallimastix frontalis; Nc N. cameroonii; Ad Aestipascuomyces dupliciliberans, Gc Ghazellomyces constrictus; Fa Feramyces austinii; Cc Caecomyces communis; Ca Cyllamyces aberensis; Am Anaeromyces mucronatus; Ac Anaeromyces concortus; Oj Orpinomyces joyonii; Pr Pecaromyces ruminatium; Ja Joblinomyces apicalis; Tm Tahromyces munnarensis; Ps Piromyces* spp*.; Be Buwchfawromyces eastonii; Kr Khoyollomyces ramosus; Cf Capellomyces forminis; Cs Capellomyces elongis; Liebetanzomyces polymorphus; Al Agrisomyces longus; Ap Aklioshbomyces papillarum; Tg Testudinimyces gracilis; Ad Astrotestudinimyces divisus.* **(C)** Experimental visualisation of RFLP profiles for selection of AGF genera and species obtained after digest of NL1-NL4 PCR product with AluI, and **(D)** after digest with Hpy188I. G3, *Feramyces* sp isolate G3; G5, *F.* sp G5; N9, *Neocallimastix cameroonii* N9; N13, *N. cameroonii* N13; *Nf*, *Neocallimastix frontalis* CoB3; *Cc*, *Caecomyces communis* SHB; *Cs*, *Capellomyces forminis*. Cap2A; *Ps*, *Piromyces sp*. SHC.

We experimentally verified the LSU-based PCR-RFLP strategy, testing the profiling of a range of AGF species: the *Feramyces* G3 and G5 and *N. cameroonii* N9 and N13 isolates described in this study, as well as *N. frontalis* CoB3 (Shen et al., 2026), *Caecomyces communis* SHB (Shen et al., 2026), *Capellomyces forminis* Cap2A (Hanafy et al., 2023), *Piromyces edwardsiae* isolate SHC (Shen et al., 2026). The DNA fragment profiles generated after digestion with AluI (Fig. 5C) and Hpy188I (Fig. 5D) enabled straightforward discrimination between these species, and, when compared to predicted fragment sizes (Table S2), led to unambiguous identification of the species based on PCR-RFLP profile alone in almost all cases. The identification of the Cap2A profile solely from fragment sizes determined from the RFLP gel would be uncertain as, due to the 20-30 bp error margin, possible matches include both *Piromyces* or *Capellomyces*. However, as visible on the gel (Fig 5A, B), including a reference sample for each of the strains allows clear distinction between the two genera. Overall, this highlights that PCR-RFLP based on NL1 and NL4 amplification and digest with AluI and Hpy188I allows excellent discrimination of AGF genera cultured so far.

### Supplementing enrichment media with vitamins enabled Khoyollomyces growth

As *Khoyollomyces* dominated AGF in zebra faecal samples (Fig. 2), but initial AGF enrichment cultures were unsuccessful, we reviewed reported growth conditions for *Khoyollomyces* (Hanafy, Lanjekar, et al., 2020; Jones et al., 2024; Schulz et al., 2025). A major difference with the medium used here was inclusion of vitamin and trace element solutions. We therefore hypothesized that the enrichment medium was lacking vitamins or trace elements required for *Khoyollomyces* growth. Indeed, addition of vitamin solution, but not trace element solution, enabled successful enrichments. The subsequently obtained isolate Z3 produced small colonies on solid medium and grain-like biomass morphology in liquid cultures of medium C (Fig. 6A). Microscopy revealed filamentous mycelia with thalli with often multiple sporangia (Fig. 6B), and monoflagellated zoospores (Fig. 6C), all consistent with characteristics of *Khoyollomyces ramosus*. The isolate’s PCR-RFLP pattern (Fig 6D) was consistent with that predicted for *Khoyollomyces* (Table S2, Fig. 5A, B), and amplicon sequencing using AGF-envS primers confirmed this identification, with 100% of reads having 99.7-100% identity to *Khoyollomyces ramosus* sequences. Notably, vitamin supplementation was required only during establishment of the *K. ramosus* culture, the isolate has been in stable cultivation in medium C for ∼ 4 months without additional vitamin supplementation.

**Figure 6.**
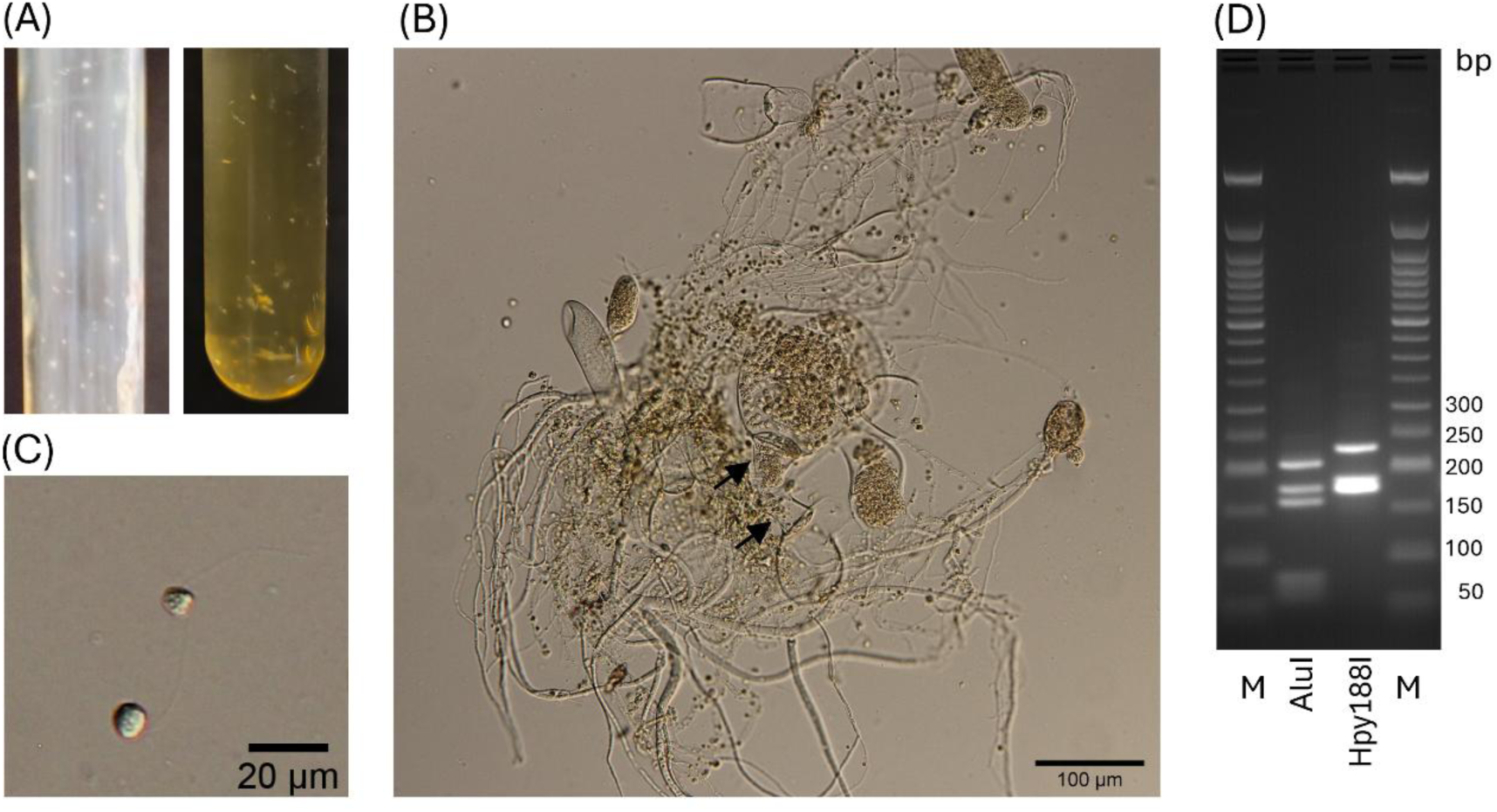
Characteristics of *Khoyollomyces* isolate Z3. **(A)** Macromorphology of isolate Z3, isolated from Zebra stool, on solid media C with glucose as carbon source (left) and on liquid media (right), **(B)** filamentous morphology as observed via microscopy, with thalli with multiple sporangia as characteristic of *Khoyollomyces* (arrow), **(C)** mono-flagellated zoospore, **(D)** LSU-based PCR-RFLP profile of isolate Z3, after digestion with AluI and Hpy118I.

### Isolation of chloramphenicol-sensitive NY08 representative from nyala

As nyala samples were rich in so far uncultured *Piromyces*-associated NY08, and considering 25% of nyala-derived isolates were previously lost when adding chloramphenicol, we repeated enrichment and isolation avoiding chloramphenicol. All 5 obtained isolates had identical PCR-RFLP profiles that were consistent with those predicted and observed for *Piromyces* (Fig. 7A, B), and Sanger sequencing of isolate N2.1 and N3.1 ITS and LSU regions confirmed NY08 representatives were obtained. Intra-isolate sequence variation was 0-0.38% across 10 LSU sequences and 0% across 3 ITS sequences. LSU sequences were 99.6-99.9% identical to sequences forming the basis for initially proposing NY08 (e.g. OP253938), indicating these belong to the same species, which we will refer to here as *Piromyces* sp. N2.1 while awaiting reclassification based on ongoing phylogenetic analysis of the *Piromycetaceae*. For both isolates of the species obtained here, small brown colonies formed on solid medium, in liquid medium mature mycelium adhered tightly to culture tubes (Fig. 7C). The filamentous hyphae were thin and wrinkly (Fig. 7D), and zoospores (8.0 ± 0.9 µm diameter, n=30) were monoflagellated, consistent with what was previously described (Hanafy et al., 2022) for *Piromyces* species.

**Figure 7.**
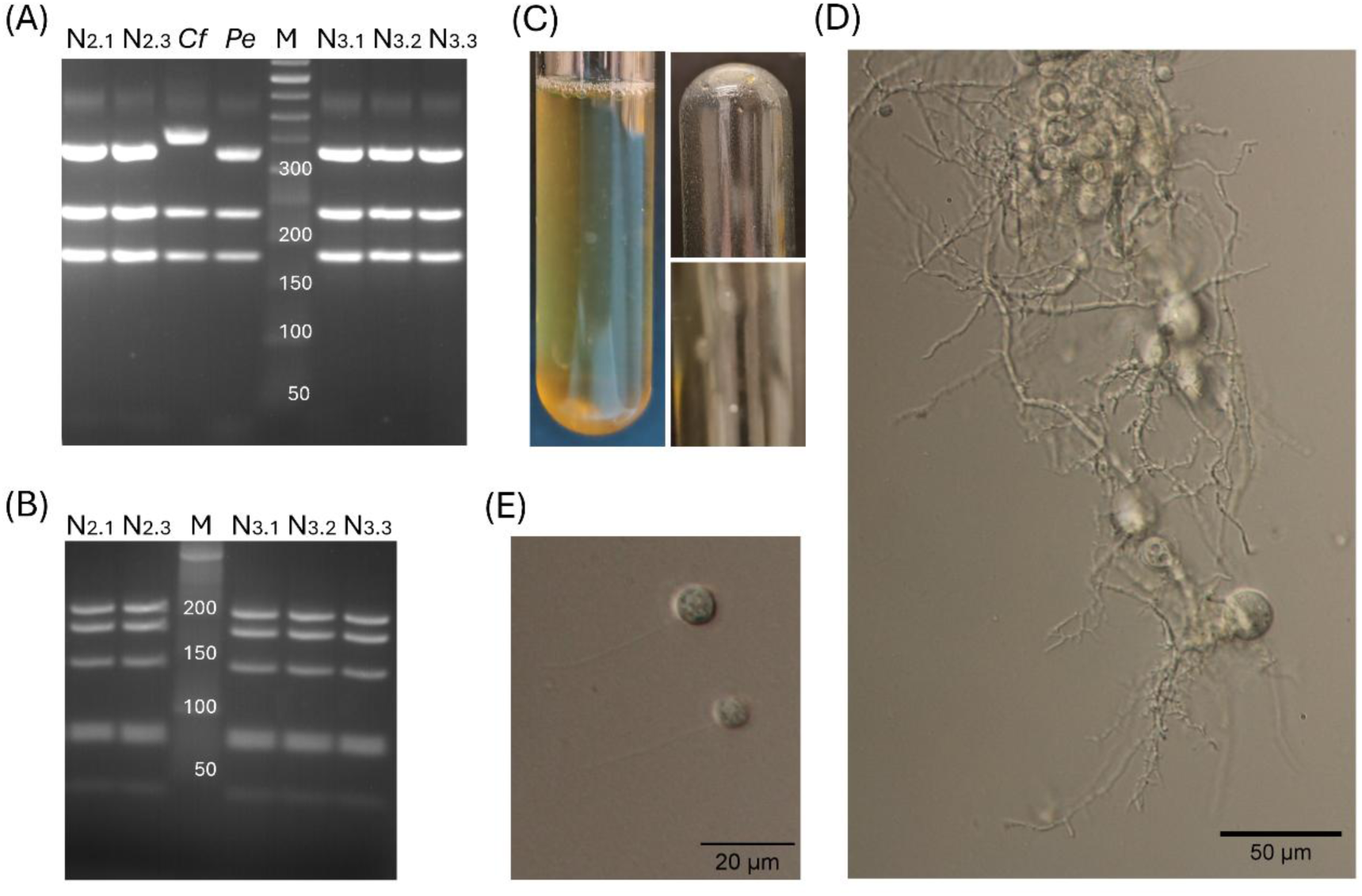
Characteristics of *Piromyces* sp isolate 2.1, a NY08 representative. **(A)** LSU-based PCR-RFLP profile of isolates N2.1, N2.3 and N3.1-N3.3, after digestion with Hpy118I, compared to the profile obtained for *Capellomyces forminis* (*Cf*) and *Piromyces edwardsiae (Pe)* **(B)** PCR-RFLP after digestion with AluI. **(C)** Macromorphology of isolate N2.1 on liquid media C with glucose as carbon source (left culture as in media, top right biofilm imaged without media), and on solid media (bottom right), **(D)** filamentous morphology as observed via microscopy, with thin, short and heavily branched filamentous rhizoids, **(E)** mono-flagellated zoospores.

## Discussion

Here we described the isolation and basic characterisation of anaerobic gut fungi representing four different genera relatively abundant in faecal samples of their host, including the so far uncultured *Piromyces* sp. N2.1 (previously proposed as novel genus NY08 (Meili et al., 2023)). We demonstrated how an LSU-based PCR-RFLP strategy that we developed and validated here can be employed to quickly and efficiently profile and identify fungal isolates. We highlight how enrichment and isolation conditions, including use of antibiotics, may be critical to isolation success.

The AGF community in the faecal samples of zebra was dominated by *Khoyollomyces,* which is line with the reported strong association of the genus *Khoyollomyces* with horses and zebras (Hanafy, Johnson, et al., 2020; Meili et al., 2023) and reported isolation of this genus from horse and zebra (Hanafy, Lanjekar, et al., 2020; Stabel et al., 2021). *Feramyces* was abundant in giraffe faecal samples, similar as described (Liggenstoffer et al., 2010; Meili et al., 2023) but isolation of this genus has only been reported from wild Barbary sheep and fallow deer (Hanafy et al., 2018). In contrast, isolation of *Neocallimastix*, which here was relatively low in abundance in the giraffe AGF community (Meili et al., 2023), has been reported (isolate GF-Ma3-1, genome available on mycocosm, isolate G341 (Stabel et al., 2021)). We report here to our knowledge for the first time, the composition of the AGF community of the nyala, a spiral-horned antelope related to the kudu, bushbuck, and eland. Results indicated low AGF diversity with two dominant species, and a high relative abundance of an uncultured fungus. We were able to capitalise on this via successful isolation of both species, which included the NY08 representative *Piromyces* sp. N2.1. Notably the phylogeny of *Piromyces* and the wider *Piromycetaceae* family is complex (Hanafy et al., 2023) and isolation of NY08 followed by obtaining genomic or transcriptomic data would aid required revision of the family’s phylogeny and clarify the phylogenetic placement of NY08.

In our workflow we obtained fungal isolates of only one species per enrichment, whereby simultaneously performed replicate enrichments yielded similar results. Others (Hanafy, Lanjekar, et al., 2020; Peng et al., 2021; Stabel et al., 2021) have reported similar outcomes. As also suggested by (Stabel et al., 2021), it seems probable that obtained isolates would be the dominant species of fungus in the faecal sample, or well adapted to the enrichment conditions. Indeed (Hanafy, Johnson, et al., 2020), who isolated one to three genera per faecal sample in a large isolation effort, reported a positive correlation between frequency of isolation of monocentric taxa (such as *Feramyces* and *Neocallimastix*) and abundance in a sample and a negative correlation of frequency of isolation with community evenness. AGF community profiling in a study including samples from 5 animal species (Joshi et al., 2022) suggested that enrichment cultures are dominated (>90% of relative abundance) by one or two genera even if up to three to four genera are each present at >10% relative abundance in the corresponding faecal sample. Notably, genera dominating enrichments did not always correspond with the most abundant genera in the faecal sample (Joshi et al., 2022), underlining how species that are well adapted or competitive under the enrichment conditions were obtained. Similarly, in our results, successful enrichment and isolation of *Khoyollomyces* Z3 required addition of vitamin solution, and for cultivation of *Piromyces* sp. N2.1 the omission of chloramphenicol, routinely used for AGF enrichments, from the cultivation medium was required. Combined, these findings suggests that enrichment cultures may be often dominated by one or few AGF genera, and isolating 15-20 colonies per enrichment, as performed here, would be more than sufficient to enable isolation of the expected one to three enriched fungal genera. It is likely that stratifying enrichment conditions (carbon source, media composition, antibiotics employed) or inoculum treatment (storage time, method) may be effective when aiming to obtain distinct genera from a sample.

We tested a workflow for the identification of fungal isolates that integrated the assessment of morphology, carbon source usage as well as molecular profiling of genetic diversity. Observations of morphology (filamentous or bulbous) and zoospore flagellation were informative in assessing diversity of fungal isolates, but as reviewed (Hanafy et al., 2022) were, in isolation non-conclusive in confirming identity. Integration of morphological analysis with molecular tools has enabled identification and description of a range of recent genera (Hanafy et al., 2017, 2022). However, in this study cloning and sequencing of ITS or LSU regions of all isolates was cost prohibitive and time consuming, prompting us to develop a PCR-RFLP strategy that proved efficient and effective in profiling individual isolates, and provided unambiguous results even in the hands of inexperienced students. With the LSU-based method described here we were able to process ∼ 20 isolates (6 ml cultures each) in 2 days for ∼ £90. The profiling of carbon source usage proved resource intensive as well as error prone and did not bring benefit here, it is best reserved for profiling selected axenic fungal isolates.

In developing the PCR-RFLP method, we capitalised on the availability of rRNA gene sequences for distinct AGF genera and species as available in a database via the AGF network website. Initially we used *in silico* analysis to generate a table of fragment profiles that can be expected when ITS-based RFLP is executed as described by (Fliegerová et al., 2006). However, the resulting multiple dominant RFLP profiles per genus made identification onerous. We aimed to improve the PCR-RFLP method by using the D1-D2 region of the 28S LSU rRNA gene sequence as marker. As AGF have considerable divergence in ITS sequence length and sequence composition, even within one species, the ITS region is not considered a good phylogenetic marker region for AGF (Edwards et al., 2017; Hanafy et al., 2023; Hanafy, Johnson, et al., 2020), and alternative approaches using the D1-D2 LSU region have been developed (Edwards et al., 2017; Hanafy, Johnson, et al., 2020; Young et al., 2022). Aligned with this, and encouraged by previous reported PCR-RFLP-based distinction of *Orpinomyces joyonii* and *Orpinomyces intercalaris* (Dagar et al., 2011) and further experiment discrimination between 6 AGF genera using this region (Dagar et al., 2014), we explored use of LSU-based PCR-RFLP. Our *in silico* prediction combined with experimental validation on axenic cultures demonstrated that profiling with a combination of Hpy118I and AluI provided discriminatory patterns across the board for most reported cultured genera. Further testing on mixed AGF communities from enrichments suggested that this method may be employed to perform initial screening of AGF genera enriched under different conditions. Such an initial screening could be performed within the timeframe of one fungal cultivation cycle (2-4 days) and could complement initial sequencing and selection of faecal samples, which is reported to be highly beneficial in directing isolation efforts for obtaining novel AGF species (Hanafy, Johnson, et al., 2020). With the ongoing expansion of the LSU database, more full-length DNA sequences are expected to become available, which will further increase the accuracy of the fragment size predictions.

## Acknowledgements

We thank Philip Mathis from the Royal Zoological Society of Scotland and the team of keepers at Edinburgh Zoo for collecting and providing faecal samples. We also thank Noha Youssef and Julia Vinzelj for providing *Capellomyces* Cap2A.

## Data Availability

Data are available in the supplementary material of this article. Sanger sequences of ITS and LSU clones of isolates are deposited in NCBI Genbank database under accession numbers PZ440954-PZ440957 and PZ441541-PZ441552, respectively. Amplicon sequencing data is available in NCBI Sequence Read Archive (SRA) under BioProject PRJNA1469515. AGF-envS-based amplicon sequencing data is available under BioProject PRJNA1469551.

---

**Figure S1.**
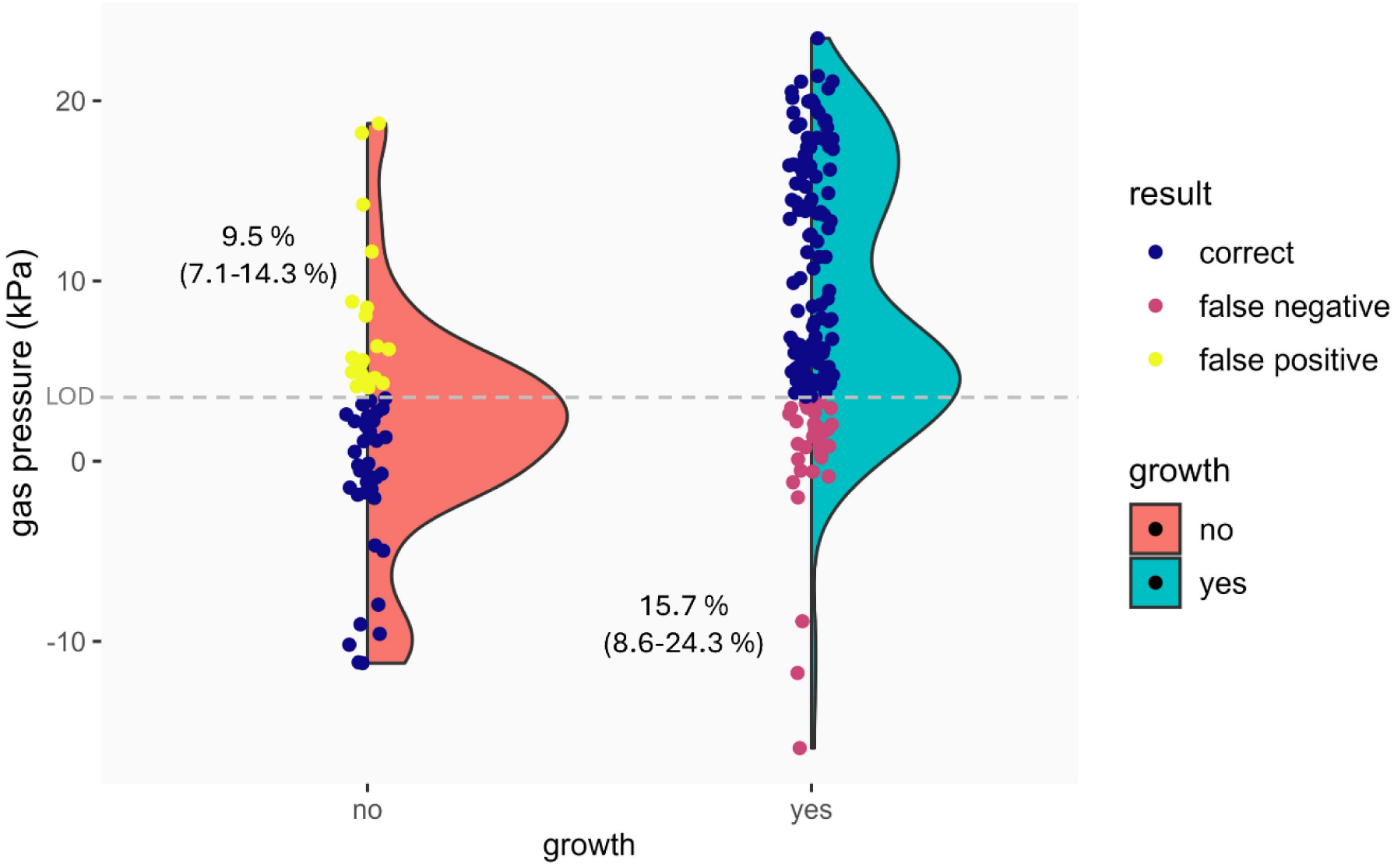
Gas pressure and visual growth observations for initial isolates from nyala and giraffe and found discrepancies between both observations.

**Figure S2.**
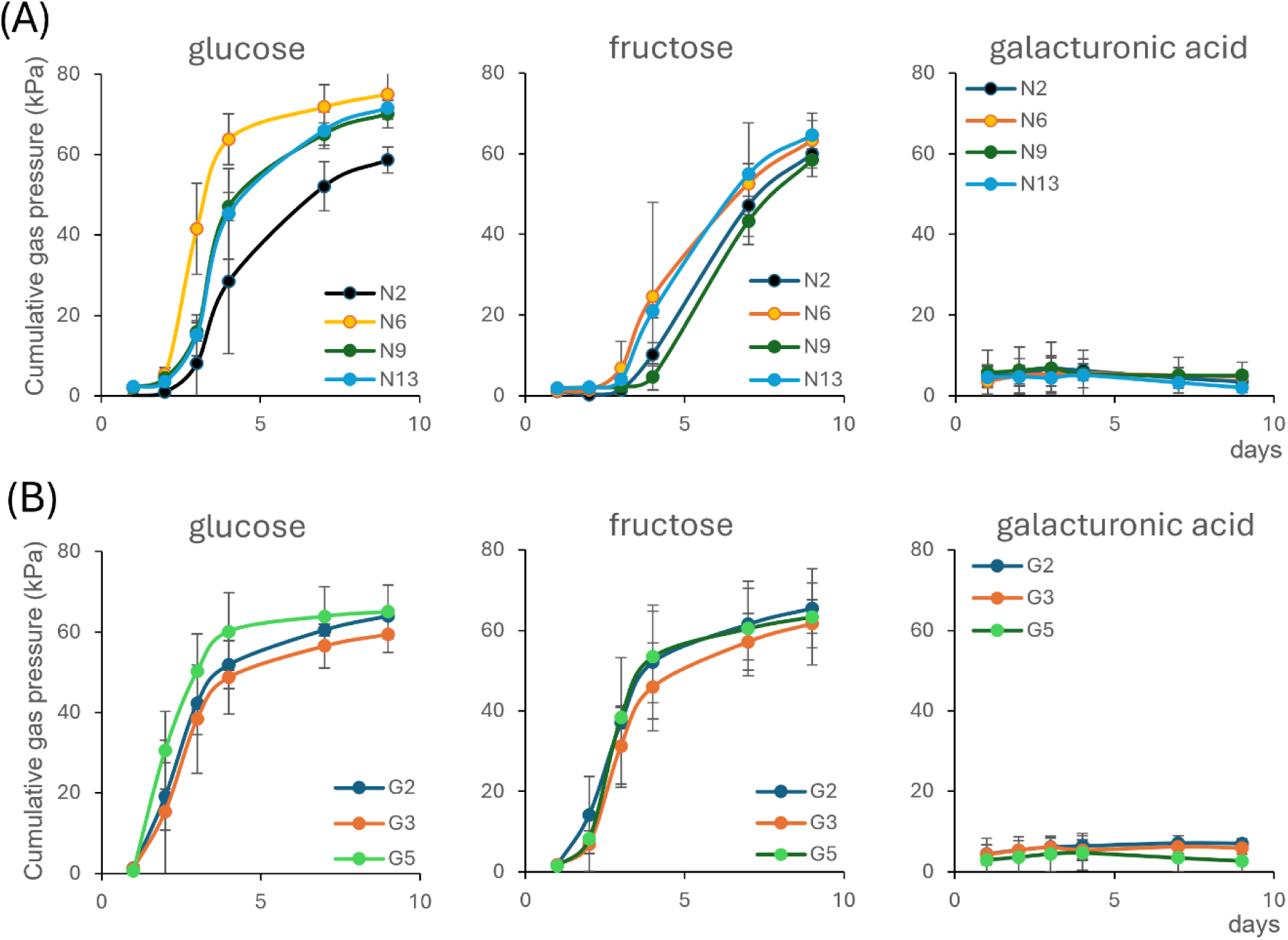
Growth curves of axenic fungal cultures. Cumulative fermentation gas production over time in cultures with glucose, fructose and galacturonic acid as sole carbon source, of **(A)** *N. cameroonii* isolates N2, N6, N9, N13 and **(B)** *F. austinii* isolates G2, G3, G5. Values are mean ± standard deviation (n=3).

**Figure S3.**
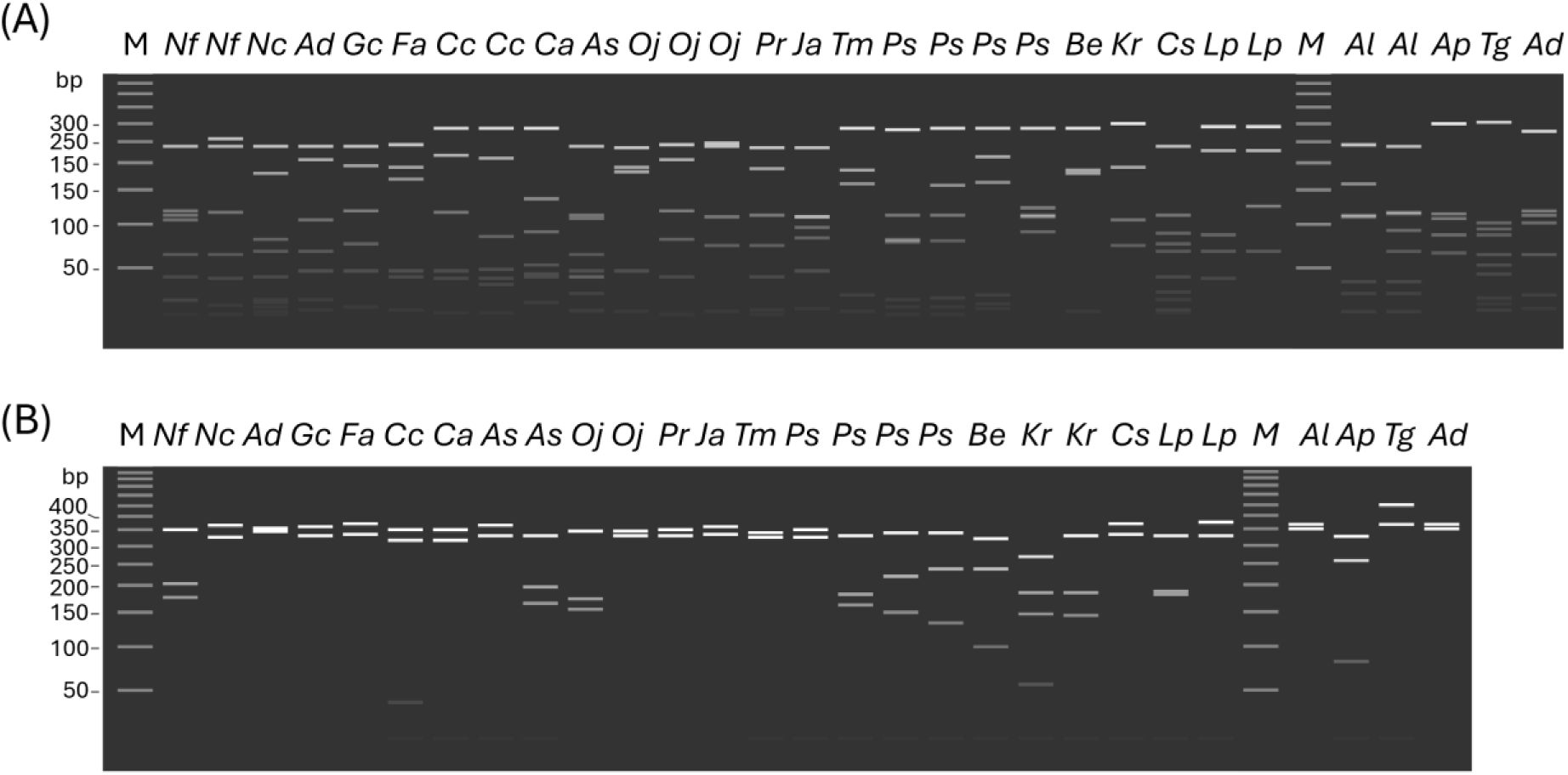
Integration of RFLP with ITS sequence information. Visualisation of RFLP profiles predicted for currently established AGF genera as derived from in silico digestion with **(A)** DraI and **(B)** HinfI digestion. *Nf Neocallimastix frontalis; Nc N. cameroonii; Ad Aestipascuomyces dupliciliberans, Gc Ghazellomyces constrictus; Fa Feramyces austinii ; Cc Caecomyces communis; Ca Cyllamyces aberensis; As Anaeromyces spp; Oj Orpinomyces joyonii, Pr Pecaromyces ruminatium; Ja Joblinomyces apicalis; Tm Tahromyces munnarensis; Ps Piromyces* spp*.; Be Buwchfawromyces eastonii; Kr Khoyollomyces ramosus; Cs Capellomyces spp.; Liebetanzomyces polymorphus; Al Agrisomyces longus; Ap Aklioshbomyces papillarum; Tg Testudinimyces gracilis; Ad Astrotestudinimyces divisus*.

**Figure S4.**
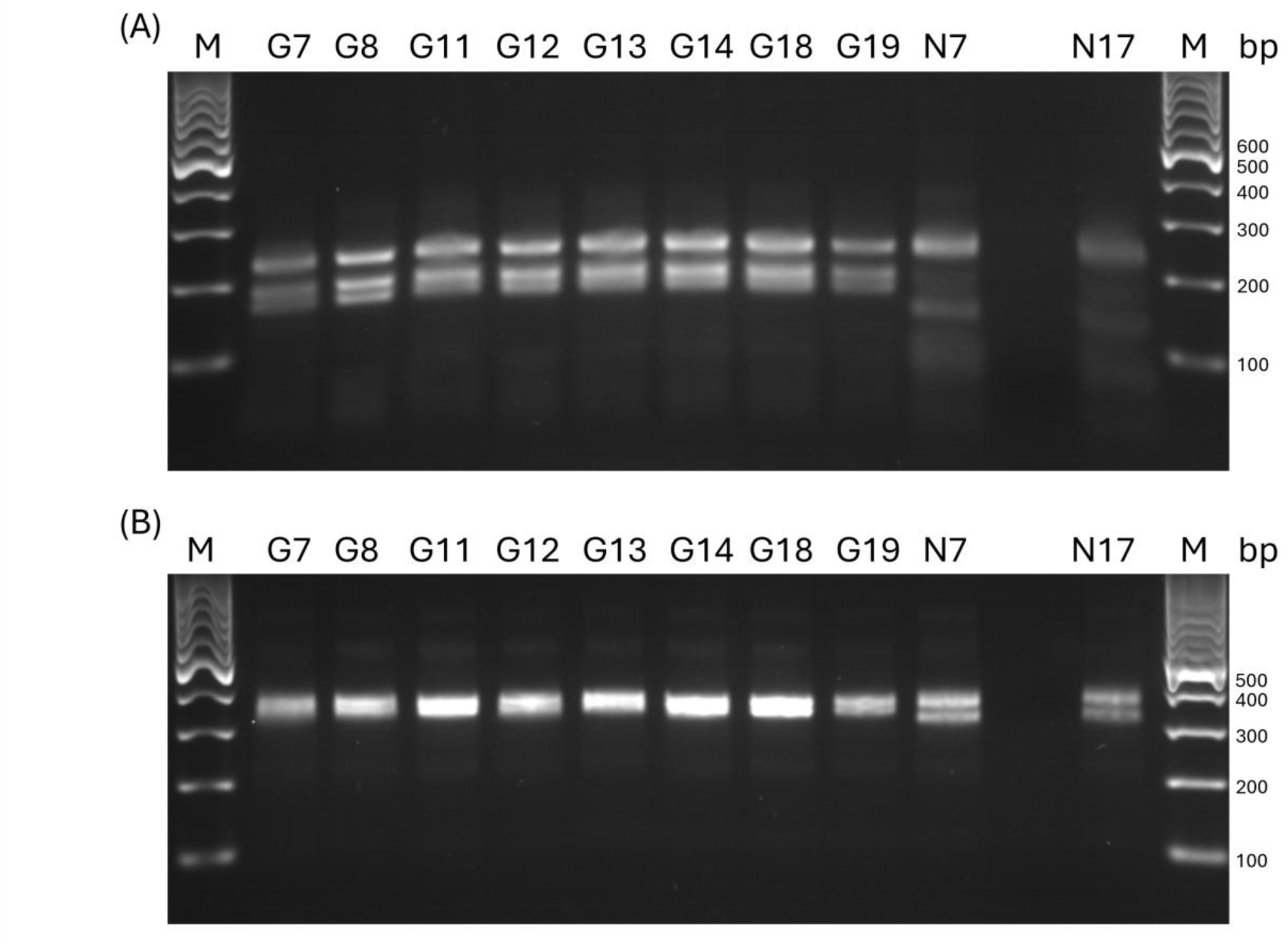
PCR-RFLP of selected isolates from giraffe. ITS rRNA region digested with **(A)** DraI or **(B)** with HinfI.

**Table S1.** ITS-based PCR-RFLP fragment predictions for cultured genera

**Table S2.** LSU-based PCR-RFLP fragment predictions for cultured genera

## References

Akin, D. E., & Borneman, W. S. (1990). Role of Rumen Fungi in Fiber Degradation. In Journal of Dairy Science (Vol. 73). 10.3168/jds.S0022-0302(90)78989-8

Dagar, S. S., Kumar, S., Griffith, G. W., Edwards, J. E., Callaghan, T. M., Singh, R., Nagpal, A. K., & Puniya, A. K. (2015). A new anaerobic fungus (Oontomyces anksri gen. nov., sp. nov.) from the digestive tract of the Indian camel (Camelus dromedarius). Fungal Biology, 119(8), 731–737. 10.1016/j.funbio.2015.04.005

Dagar, S. S., Kumar, S., Mudgil, P., Singh, R., & Puniya, A. K. (2011). D1/D2 domain of large-subunit ribosomal DNA for differentiation of Orpinomyces spp. Applied and Environmental Microbiology, 77(18), 6722–6725. 10.1128/AEM.05441-11

Dagar, S. S., Kumar, S., Pitta D. W, Edwards, J., Callaghan, T., Griffith, G., Mudgil, P., & Puniya, A. (2014). Large-subunit rDNA based differentiation of anaerobic rumen fungi using restriction fragment length polymorphism. Linking Animal Science and Animal Agriculture: Meeting the Global Demands of 2050 92, ADSA-ASAS-CSAS Joint Annual Meeting, 340–340. https://www.adsa.org/Portals/0/SiteContent/Docs/Meetings/PastMeetings/Annual/2014/JAM14-Abstracts.pdf

Davies, D. R., Theodorou, M. K., Lawrence, M. I. G., & Trinci, A. P. J. (1993). Distribution of anaerobic fungi in the digestive tract of cattle and their survival in faeces. Journal of General Microbiology, 139(6), 1395–1400. 10.1099/00221287-139-6-1395

Edwards, J. E., Forster, R. J., Callaghan, T. M., Dollhofer, V., Dagar, S. S., Cheng, Y., Chang, J., Kittelmann, S., Fliegerova, K., Puniya, A. K., Henske, J. K., Gilmore, S. P., O’Malley, M. A., Griffith, G. W., & Smidt, H. (2017). PCR and Omics Based Techniques to Study the Diversity, Ecology and Biology of Anaerobic Fungi: Insights, Challenges and Opportunities. Frontiers in Microbiology | Www.Frontiersin.Org, 8, 1657. 10.3389/fmicb.2017.01657

Elshahed, M. S., Hanafy, R. A., Cheng, Y., Dagar, S. S., Edwards, J. E., Flad, V., Fliegerová, K. O., Griffith, G. W., Kittelmann, S., Lebuhn, M., O’malley, M. A., Podmirseg, S. M., Solomon, K. V., Vinzelj, J., Young, D., & Youssef, N. H. (2022). Characterization and rank assignment criteria for the anaerobic fungi (Neocallimastigomycota). International Journal of Systematic and Evolutionary Microbiology, 72(7), 005449. 10.1099/IJSEM.0.005449/CITE/REFWORKS

Fliegerová, K., Mrázek, J., & Voigt, K. (2006). Differentiation of anaerobic polycentric fungi by rDNA PCR-RFLP. Folia Microbiologica, 51(4), 273–277. 10.1007/BF02931811

Griffith, G. W., Ozkose, E., Theodorou, M. K., & Davies, D. R. (2009). Diversity of anaerobic fungal populations in cattle revealed by selective enrichment culture using different carbon sources. Fungal Ecology, 2(2), 87–97. 10.1016/j.funeco.2009.01.005

Gruninger, R. J., Puniya, A. K., Callaghan, T. M., Edwards, J. E., Youssef, N., Dagar, S. S., Fliegerova, K., Griffith, G. W., Forster, R., Tsang, A., McAllister, T., & Elshahed, M. S. (2014). Anaerobic fungi (phylum Neocallimastigomycota): advances in understanding their taxonomy, life cycle, ecology, role and biotechnological potential. FEMS Microbiology Ecology, 90(1), 1–17. 10.1111/1574-6941.12383

Haitjema, C. H., Gilmore, S. P., Henske, J. K., Solomon, K. V., De Groot, R., Kuo, A., Mondo, S. J., Salamov, A. A., LaButti, K., Zhao, Z., Chiniquy, J., Barry, K., Brewer, H. M., Purvine, S. O., Wright, A. T., Hainaut, M., Boxma, B., Van Alen, T., Hackstein, J. H. P., … O’Malley, M. A. (2017). A parts list for fungal cellulosomes revealed by comparative genomics. Nature Microbiology, 2. 10.1038/nmicrobiol.2017.87

Hanafy, R. A., Dagar, S. S., Griffith, G. W., Pratt, C. J., Youssef, N. H., & Elshahed, M. S. (2022). Taxonomy of the anaerobic gut fungi (Neocallimastigomycota): a review of classification criteria and description of current taxa. International Journal of Systematic and Evolutionary Microbiology, 72(7). 10.1099/ijsem.0.005322

Hanafy, R. A., Elshahed, M. S., Liggenstoffer, A. S., Griffith, G. W., & Youssef, N. H. (2017). Pecoramyces ruminantium, gen. nov., sp. nov., an anaerobic gut fungus from the feces of cattle and sheep. Mycologia, 109(2), 231–243. 10.1080/00275514.2017.1317190

Hanafy, R. A., Elshahed, M. S., & Youssef, N. H. (2018). Feramyces austinii, gen. Nov., sp. nov., an anaerobic gut fungus from rumen and fecal samples of wild barbary sheep and fallow deer. Mycologia, 110(3), 513–525. 10.1080/00275514.2018.1466610

Hanafy, R. A., Johnson, B., Youssef, N. H., & Elshahed, M. S. (2020). Assessing anaerobic gut fungal diversity in herbivores using D1/D2 large ribosomal subunit sequencing and multi-year isolation. Environmental Microbiology, 22(9), 3883–3908. 10.1111/1462-2920.15164;PAGE:STRING:ARTICLE/CHAPTER

Hanafy, R. A., Lanjekar, V. B., Dhakephalkar, P. K., Callaghan, T. M., Dagar, S. S., Griffith, G. W., Elshahed, M. S., & Youssef, N. H. (2020). Seven new Neocallimastigomycota genera from wild, zoo-housed, and domesticated herbivores greatly expand the taxonomic diversity of the phylum. Mycologia, 112(6), 1212–1239. 10.1080/00275514.2019.1696619

Hanafy, R. A., Wang, Y., Stajich, J. E., Pratt, C. J., Youssef, N. H., & Elshahed, M. S. (2023). Phylogenomic analysis of the Neocallimastigomycota: proposal of Caecomycetaceae fam. nov., Piromycetaceae fam. nov., and emended description of the families Neocallimastigaceae and Anaeromycetaceae. International Journal of Systematic and Evolutionary Microbiology, 73(2). 10.1099/IJSEM.0.005735

Hausner, G., Inglis, G. D., Yanke, L. J., Kawchuk, L. M., & McAllister, T. A. (2011). Analysis of restriction fragment length polymorphisms in the ribosomal DNA of a selection of anaerobic chytrids. Https://Doi.Org/10.1139/B00-067, 78(7), 917–927. 10.1139/b00-067

Hooker, C. A., Hanafy, R., Hillman, E. T., Muñoz Briones, J., & Solomon, K. V. (2023). A genetic engineering toolbox for the lignocellulolytic anaerobic gut fungus Neocallimastix frontalis. ACS Synthetic Biology, 12(4), 1034. 10.1021/ACSSYNBIO.2C00502

Jones, A. L., Pratt, C. J., Meili, C. H., Soo, R. M., Hugenholtz, P., Elshahed, M. S., & Youssef, N. H. (2024). Anaerobic gut fungal communities in marsupial hosts. MBio, 15(2). 10.1128/MBIO.03370-23,

Joshi, A., Young, D., Huang, L., Mosberger, L., Munk, B., Vinzelj, J., Flad, V., Sczyrba, A., Griffith, G. W., Podmirseg, S. M., Warthmann, R., Lebuhn, M., & Insam, H. (2022). Effect of Growth Media on the Diversity of Neocallimastigomycetes from Non-Rumen Habitats. Microorganisms 2022, Vol. 10, Page 1972, 10(10), 1972. 10.3390/MICROORGANISMS10101972

Kittelmann, S., Naylor, G. E., Koolaard, J. P., & Janssen, P. H. (2012). A proposed taxonomy of anaerobic fungi (class neocallimastigomycetes) suitable for large-scale sequence-based community structure analysis. PLoS ONE, 7(5). 10.1371/JOURNAL.PONE.0036866,

Kopečný, J., & Hodrová, B. (1995). Pectinolytic enzymes of anaerobic fungi. Letters in Applied Microbiology, 20(5), 312–316. 10.1111/J.1472-765X.1995.TB00453.X

Kozich, J. J., Westcott, S. L., Baxter, N. T., Highlander, S. K., & Schloss, P. D. (2013). Development of a dual-index sequencing strategy and curation pipeline for analyzing amplicon sequence data on the MiSeq Illumina sequencing platform. Applied and Environmental Microbiology, 79(17), 5112–5120. 10.1128/AEM.01043-13

Lee, S. S., Ha, J. K., & Cheng, K. J. (2000). Relative contributions of bacteria, protozoa, and fungi to in vitro degradation of orchard grass cell walls and their interactions. Applied and Environmental Microbiology, 66(9), 3807–3813. 10.1128/AEM.66.9.3807-3813.2000

Liggenstoffer, A. S., Youssef, N. H., Couger, M. B., & Elshahed, M. S. (2010). Phylogenetic diversity and community structure of anaerobic gut fungi (phylum Neocallimastigomycota) in ruminant and non-ruminant herbivores. ISME Journal, 4(10), 1225–1235. 10.1038/ISMEJ.2010.49,

Ma, Y., Li, Y., Li, Y., Cheng, Y., & Zhu, W. (2020). The enrichment of anaerobic fungi and methanogens showed higher lignocellulose degrading and methane producing ability than that of bacteria and methanogens. World Journal of Microbiology and Biotechnology, 36(9), 125. 10.1007/s11274-020-02894-3

McMurdie, P. J., & Holmes, S. (2013). phyloseq: An R Package for Reproducible Interactive Analysis and Graphics of Microbiome Census Data. PLOS ONE, 8(4), e61217. 10.1371/journal.pone.0061217

Meili, C. H., Jones, A. L., Arreola, A. X., Habel, J., Pratt, C. J., Hanafy, R. A., Wang, Y., Yassin, A. S., TagElDein, M. A., Moon, C. D., Janssen, P. H., Shrestha, M., Rajbhandari, P., Nagler, M., Vinzelj, J. M., Podmirseg, S. M., Stajich, J. E., Goetsch, A. L., Hayes, J., … Elshahed, M. S. (2023). Patterns and determinants of the global herbivorous mycobiome. Nature Communications, 14(1). 10.1038/S41467-023-39508-Z

Meili, C. H., TagElDein, M. A., Jones, A. L., Moon, C. D., Andrews, C., Kirk, M. R., Janssen, P. H., Yeoman, C. J., Grace, S., Borgogna, J. L. C., Foote, A. P., Nagy, Y. I., Kashef, M. T., Yassin, A. S., Elshahed, M. S., & Youssef, N. H. (2024). Diversity and community structure of anaerobic gut fungi in the rumen of wild and domesticated herbivores. Applied and Environmental Microbiology, 90(2). 10.1128/aem.01492-23

Mizrahi, I., Wallace, R. J., & Moraïs, S. (2021). The rumen microbiome: balancing food security and environmental impacts. Nature Reviews Microbiology 2021 19:9, 19(9), 553–566. 10.1038/s41579-021-00543-6

Nagpal, R., Puniya, A. K., Sehgal, J. P., & Singh, K. (2012). Survival of anaerobic fungus Caecomyces sp. in various preservation methods: A comparative study. Mycoscience, 53(6), 427–432. 10.1007/S10267-012-0187-Y/FIGURES/2

Nicholson, M. J., McSweeney, C. S., Mackie, R. I., Brookman, J. L., & Theodorou, M. K. (2010). Diversity of anaerobic gut fungal populations analysed using ribosomal ITS1 sequences in faeces of wild and domesticated herbivores. Anaerobe, 16(2), 66–73. 10.1016/j.anaerobe.2009.05.003

Oksanen, J., Simpson, G. L., Blanchet, F. G., Kindt, R., Legendre, P., Minchin, P. R., O’Hara, R. B., Solymos, P., Stevens, M. H. H., Szoecs, E., Wagner, H., Barbour, M., Bedward, M., Bolker, B., Borcard, D., Borman, T., Carvalho, G., Chirico, M., De Caceres, M., … Weedon, J. (2026). vegan: Community Ecology Package. R package version 2.8-0. In CRAN: Contributed Packages. https://vegandevs.github.io/vegan/

Pagès, H., Aboyoun, P., Gentleman, R., & DebRoy, S. (2025). Biostrings: Efficient manipulation of biological strings.. https://bioconductor-source.r-universe.dev/Biostrings/citation

Peng, X., Wilken, S. E., Lankiewicz, T. S., Gilmore, S. P., Brown, J. L., Henske, J. K., Swift, C. L., Salamov, A., Barry, K., Grigoriev, I. V., Theodorou, M. K., Valentine, D. L., & O’Malley, M. A. (2021). Genomic and functional analyses of fungal and bacterial consortia that enable lignocellulose breakdown in goat gut microbiomes. Nature Microbiology, 6(4), 499–511. 10.1038/s41564-020-00861-0

Pratt, C. J., Meili, C. H., Jones, A. L., Jackson, D. K., England, E. E., Wang, Y., Hartson, S., Rogers, J., Elshahed, M. S., & Youssef, N. H. (2024). Anaerobic fungi in the tortoise alimentary tract illuminate early stages of host-fungal symbiosis and Neocallimastigomycota evolution. Nature Communications, 15(1). 10.1038/s41467-024-47047-4

Schloss, P. D., Westcott, S. L., Ryabin, T., Hall, J. R., Hartmann, M., Hollister, E. B., Lesniewski, R. A., Oakley, B. B., Parks, D. H., Robinson, C. J., Sahl, J. W., Stres, B., Thallinger, G. G., Van Horn, D. J., & Weber, C. F. (2009). Introducing mothur: open-source, platform-independent, community-supported software for describing and comparing microbial communities. Applied and Environmental Microbiology, 75(23), 7537–7541. 10.1128/AEM.01541-09

Schulz, K. E., Scholz, D., Sikirić, A. R., Rambow, D., Neumann, A., & Ochsenreither, K. (2025). Anaerobic gut fungi as biocatalysts: metabolic and physiological analysis of anaerobic gut fungi under diverse cultivation conditions. Frontiers in Microbiology, 16. 10.3389/FMICB.2025.1662047/PDF

Seshadri, R., Leahy, S. C., Attwood, G. T., Teh, K. H., Lambie, S. C., Cookson, A. L., Eloe-Fadrosh, E. A., Pavlopoulos, G. A., Hadjithomas, M., Varghese, N. J., Paez-Espino, D., Palevich, N., Janssen, P. H., Ronimus, R. S., Noel, S., Soni, P., Reilly, K., Atherly, T., Ziemer, C., … Kelly, W. J. (2018). Cultivation and sequencing of rumen microbiome members from the Hungate1000 Collection. Nature Biotechnology, 36(4), 359–367. 10.1038/nbt.4110

Shen, S., Matthews, J. L., Li, S., & van Munster, J. M. (2026). Anaerobic fungi Caecomyces communis, Neocallimastix frontalis and Piromyces edwardsiae sp. nov. have distinct effects on plant fibres during digestion. Royal Society Open Science, 13(3). 10.1098/RSOS.251684

Sievers, F., Wilm, A., Dineen, D., Gibson, T. J., Karplus, K., Li, W., Lopez, R., McWilliam, H., Remmert, M., Söding, J., Thompson, J. D., & Higgins, D. G. (2011). Fast, scalable generation of high-quality protein multiple sequence alignments using Clustal Omega. Molecular Systems Biology, 7, 539. 10.1038/MSB.2011.75

Solden, L. M., Naas, A. E., Roux, S., Daly, R. A., Collins, W. B., Nicora, C. D., Purvine, S. O., Hoyt, D. W., Schückel, J., Jørgensen, B., Willats, W., Spalinger, D. E., Firkins, J. L., Lipton, M. S., Sullivan, M. B., Pope, P. B., & Wrighton, K. C. (2018). Interspecies cross-feeding orchestrates carbon degradation in the rumen ecosystem. Nature Microbiology, 3(11), 1274–1284. 10.1038/s41564-018-0225-4

Solomon, K. V, Haitjema, C. H., Henske, J. K., Gilmore, S. P., Borges-Rivera, D., Lipzen, A., Brewer, H. M., Purvine, S. O., Wright, A. T., Theodorou, M. K., Grigoriev, I. V, Regev, A., Thompson, D. A., & O’Malley, M. A. (2016). Early-branching gut fungi possess a large, comprehensive array of biomass-degrading enzymes. Science., 351(6278), 1192–1195. 10.1126/science.aad1431

Stabel, M., Hanafy, R. A., Schweitzer, T., Greif, M., Aliyu, H., Flad, V., Young, D., Lebuhn, M., Elshahed, M. S., Ochsenreither, K., & Youssef, N. H. (2020). Aestipascuomyces dupliciliberans gen. Nov, sp. nov., the first cultured representative of the uncultured SK4 clade from aoudad sheep and alpaca. Microorganisms, 8(11), 1–17. 10.3390/microorganisms8111734

Stabel, M., Schweitzer, T., Haack, K., Gorenflo, P., Aliyu, H., & Ochsenreither, K. (2021). Isolation and Biochemical Characterization of Six Anaerobic Fungal Strains from Zoo Animal Feces. Microorganisms, 9(8). 10.3390/MICROORGANISMS9081655

Teunissen, M. J., Op Den Camp, H. J. M., Orpin, C. G., Huis In ’T Veld, J. H. J., & Vogels, G. D. (1991). Comparison of growth characteristics of anaerobic fungi isolated from ruminant and non-ruminant herbivores during cultivation in a defined medium. Journal of General Microbiology, 137(6), 1401–1408. 10.1099/00221287-137-6-1401

Theodorou, M. K., Brookman, J., & Trinci, A. P. J. (2005). Anaerobic fungi. Methods in Gut Microbial Ecology for Ruminants, 55–66. 10.1007/1-4020-3791-0_5

Theodorou, M. K., Davies, D. R., Nielsen, B. B., Lawrence, M. I. G., & Trinci, A. P. J. (1995). Determination of growth of anaerobic fungi on soluble and cellulosic substrates using a pressure transducer. Microbiology, 141(3), 671–678. 10.1099/13500872-141-3-671

Trinci, A. P. J., Davies, D. R., Gulli, K., Lawrence!, M. I., Nielsew, B. B., Rickers, A., And, I., & Theodorou, M. K. (1994). Anaerobic fungi in herbivorous animals. Mycol. Res, 98(2), 129–152. 10.1016/S0953-7562(09)80178-0

Vinzelj, J., Joshi, A., Young, D., Begovic, L., Peer, N., Mosberger, L., Luedi, K. C. S., Insam, H., Flad, V., Nagler, M., & Podmirseg, S. M. (2022). No time to die: Comparative study on preservation protocols for anaerobic fungi. Frontiers in Microbiology, 13, 978028. 10.3389/FMICB.2022.978028/BIBTEX

Vinzelj, J., Nash, K., Jones, A. L., Young, R. T., Meili, C. H., Pratt, C. J., Wang, Y., Elshahed, M. S., & Youssef, N. H. (2025). The anaerobic gut fungal community in ostriches (Struthio camelus). ISME Communications. 10.1093/ISMECO/YCAF144

Wang, X., Liu, X., & Groenewald, J. Z. (2017). Phylogeny of anaerobic fungi (phylum Neocallimastigomycota), with contributions from yak in China. Antonie van Leeuwenhoek, International Journal of General and Molecular Microbiology, 110(1), 87–103. 10.1007/s10482-016-0779-1

Wilken, St. E., Leggieri, P. A., Kerdman-Andrade, C., Reilly, M., Theodorou, M. K., & O’Malley, M. A. (2020). An Arduino based automatic pressure evaluation system to quantify growth of non-model anaerobes in culture. AIChE Journal, 66(12), e16540. 10.1002/aic.16540

Wright, E. S. (2024). Fast and Flexible Search for Homologous Biological Sequences with DECIPHER v3. The R Journal, 16(2), 191–200. https://www.bioconductor.org/packages/release/bioc/html/DECIPHER.html

Young, D., Joshi, A., Huang, L., Munk, B., Wurzbacher, C., Youssef, N. H., Elshahed, M. S., Moon, C. D., Ochsenreither, K., Griffith, G. W., Callaghan, T. M., Sczyrba, A., Lebuhn, M., & Flad, V. (2022). Simultaneous Metabarcoding and Quantification of Neocallimastigomycetes from Environmental Samples: Insights into Community Composition and Novel Lineages. Microorganisms, 10(9), 1749. 10.3390/MICROORGANISMS10091749/S1

